# Lifelong genetically lowered sclerostin and risk of cardiovascular disease

**DOI:** 10.1101/531004

**Authors:** Jonas Bovijn, Kristi Krebs, Chia-Yen Chen, Ruth Boxall, Jenny C. Censin, Teresa Ferreira, Sara L. Pulit, Craig A. Glastonbury, Samantha Laber, Iona Y. Millwood, Kuang Lin, Liming Li, Zhengming Chen, Lili Milani, Robin G. Walters, Reedik Mägi, Benjamin M. Neale, Cecilia M. Lindgren, Michael V. Holmes

## Abstract

**Background:** Inhibition of sclerostin is a novel therapeutic approach to lowering fracture risk. However, phase III randomised controlled trials (RCTs) of romosozumab, a monoclonal antibody that inhibits sclerostin, suggest an imbalance of serious cardiovascular events.

**Methods:** We used two independent genetic variants (rs7209826 and rs188810925) in *SOST* (encoding sclerostin) associated with bone mineral density (BMD) as proxies for therapeutic inhibition of sclerostin. We estimated the effects on risk of osteoporosis, fracture, coronary heart disease (CHD) and a further 22 cardiometabolic risk factors and diseases, by combining data from up to 478,967 participants of European ancestry from three prospective cohorts and up to 1,030,836 participants from nine GWAS consortia. In addition, we performed meta-analyses of cardiovascular outcome data from phase III RCTs of romosozumab.

**Results:** Meta-analysis of RCTs identified a higher risk of cardiac ischemic events in patients randomised to romosozumab (25 events among 4,298 individuals; odds ratio [OR] 2·98; 95% confidence interval [CI], 1·18 to 7·55; P=0·017). Scaled to the equivalent dose of romosozumab (210mg/month; 0·09 g/cm^2^ higher BMD), the *SOST* variants associated with lower risk of fracture (OR, 0·59; 95% CI, 0·54-0·66; P= 1·4×10^−24^), and osteoporosis (OR, 0·43; 95% CI, 0·36-0·52; P=2·4×10^−18^). The *SOST* variants associated with higher risk of myocardial infarction and/or coronary revascularisation (69,649 cases; OR, 1·18; 95% CI, 1·06-1·32; P=0·003) and type 2 diabetes (OR 1·15; 95% CI, 1·05-1·27; P=0·003), higher systolic blood pressure (1·3mmHg; 95% CI 0·8-1·9; P=5·9×10^−6^) and waist-to-hip-ratio adjusted for BMI (0·05 SDs; 95% CI, 0·02 to 0·08; P=8·5×10^−4^).

**Conclusions:** Genetically and therapeutically lowered sclerostin leads to higher risk of cardiovascular events. Rigorous evaluation of the cardiovascular safety of romosozumab and other sclerostin inhibitors is warranted.

## Introduction

Osteoporosis, a disorder of bone demineralisation, is a common condition associated with considerable morbidity and mortality, particularly among postmenopausal women and the elderly.^1^ A range of therapeutics for osteoporosis is currently available, though these agents are plagued by poor adherence (fewer than 40% of patients prescribed oral bisphosphonates still take these medications after one year^2^), occurrence of rare but serious adverse events (e.g., bisphosphonate-induced osteonecrosis of the jaw), high cost, and uncertainty regarding long-term efficacy.^3^ There is therefore a substantial demand for effective, safe and well-tolerated anti-osteoporotic therapies.^4^

Sclerostin, a glycoprotein encoded by the *SOST* gene, is a negative regulator of bone formation that is secreted by osteocytes. It inhibits Wnt signaling, which leads to down-regulation of osteoblast development and function.^5^ Loss-of-function mutations in *SOST* lead to sclerosteosis, a rare autosomal recessive condition characterised by bone overgrowth.^6^ Similarly, van Buchem’s disease, another rare autosomal recessive condition with a clinical picture similar to sclerosteosis (albeit generally milder), is caused by a deletion of a *SOST*-specific regulatory element.^7^ The discovery of functional variants in *SOST* as the underlying cause of these rare conditions of bone overgrowth nearly 2 decades ago, led to the development of sclerostin inhibitors as a treatment for osteoporosis.^8^

Three anti-sclerostin monoclonal antibodies have been, or are, currently in clinical development,^5^ including romosozumab (Amgen, UCB) and blosozumab (Eli Lilly and Company) for osteoporosis and setrusumab (Mereo BioPharma), currently in phase IIb for osteogenesis imperfecta. Despite phase II results showing significant improvements in bone mineral density (BMD),^9^ a biomarker used to evaluate the effect of anti-osteoporotic treatments, clinical development for blosozumab was halted in 2015, reportedly due to injection site reactions.^10^ Phase II and III randomised controlled trials (RCTs) have shown that romosozumab is effective at increasing BMD in both men and women, whilst also reducing vertebral and non-vertebral fracture risk in women.^11–14^ However, adverse event data reported in the phase III BRIDGE and ARCH trials (in men and post-menopausal women, respectively) have suggested that romosozumab may be associated with an excess risk of cardiovascular events.^13,14^ Concerns about the cardiovascular safety of romosozumab, and sclerostin-inhibition more generally, have previously been raised.^8,15–17^

In January 2019, the Japanese Ministry of Health, Labor and Welfare approved the use of romosozumab in men and postmenopausal women with osteoporosis at high risk of fracture.^18^ The US Food and Drug Administration (FDA) and the European Medicines Agency (EMA) are currently considering licensing applications for romosozumab for the treatment of osteoporosis in men and postmenopausal women at increased risk of fracture.^19,20^ A detailed appraisal of the cardiovascular effects of sclerostin inhibition by exploiting randomised data from orthogonal sources is both timely and warranted, and may assist in the evaluation of whether this class of drugs represents a rational and safe therapeutic strategy for the prevention of fracture.

Naturally-occurring human genetic variation can serve as a proxy for therapeutic stimulation or inhibition of a drug target, presenting a valuable opportunity to assess the likely consequences of modifying a therapeutic target on both the intended therapeutic effects and target-mediated adverse drug reactions.^21^ The application of this approach in a Mendelian randomisation framework has previously shown to recapitulate known clinical effects of drug target modulation.^22–26^ In this study, we used this genetic approach to examine the effect of BMD-increasing alleles in the *SOST* locus (as a proxy for sclerostin inhibition) on the risk of bone fracture, osteoporosis, cardiovascular risk factors and disease, to shed light on whether treatment with sclerostin inhibitors is likely to adversely impact on cardiovascular disease.

## Methods

### Study population

We examined individual-level genotypic and phenotypic data for 502,617 subjects in UK Biobank (UKBB), a population-based cohort based in the United Kingdom. Following quality control, 423,761 subjects of white ethnicity were included for further analysis (see **Supplementary Methods** and **Table S1** in the appendix for full quality control descriptions and baseline characteristics of participants in UKBB). In addition, we included data from a further two European-ancestry cohorts: Partners HealthCare Biobank (PHB; up to 19,132 subjects) and Estonian Biobank (EGCUT; up to 36,074 subjects). Trans-ethnic replication was attempted in China Kadoorie Biobank (CKB; up to 81,546 subjects). See **Supplementary Methods** in the appendix for further detail on each cohort.

We supplemented these data with summary-level data from 9 genome-wide association study (GWAS) consortia, including data for BMD (ultrasound-derived estimated heel-bone BMD [up to 142,487 individuals] and dual-energy x-ray absorptiometry (DXA)-derived BMD [up to 49,988]), coronary heart disease (CHD; up to 60,801 cases and 123,504 controls) and myocardial infarction, stroke (up to 40,585 cases and 406,111 controls), atrial fibrillation (up to 60,620 cases and 970,216 controls), type 2 diabetes mellitus (T2D; up to 74,124 cases and 824,006 controls), glycemic traits (up to 123,665 individuals), serum lipid fractions (up to 92,804 individuals), anthropometric traits (up to 222,233 individuals), and renal function (up to 110,515 individuals). Further details on each consortium are provided in the **Supplementary Methods** and **Table S2** in the appendix. An overview of the overall study design is shown in **Figure S1** in the appendix. All studies contributing data to these analyses were approved by their local ethics committees (details in **appendix**).

### Genetic instrument selection and validation

A recent large-scale GWAS for estimated heel-bone BMD (eBMD), conducted in 142,487 individuals,^27^ identified two conditionally independent (r^2^=0·13 among European ancestry individuals in UKBB) genetic variants in the *SOST* locus associated with eBMD: rs7209826 (A>G, G-allele frequency in UKBB = 40%) and rs188810925 (G>A, A-allele frequency = 8%). Both SNPs are located ~35kb downstream from *SOST* and fall within or near a 52kb area that contains the van Buchem disease deletion (**Figure S2** in the appendix), a region previously shown to affect *SOST* expression in human bone.^7^ Recent functional evidence has also shown that SNPs in this area (one of which, rs7220711, is in high LD [r^2^=0·99] with rs7209826) regulate *SOST* expression via differential transcription factor binding.^28^ Previous Mendelian randomisation studies have also made use of non-coding variants with an effect on gene expression as proxies for pharmacologic modulation of the same target-gene.^22,26,29,30^

We extracted estimates for these two SNPs (rs7209826 and rs188810925) from GWAS consortia listed in **Table S2** in the appendix. For GWAS datasets that did not include these variants, we selected variants in high LD (r^2^>0·9) with our selected SNPs (See **Supplementary Methods** in the appendix for details). We selected rs7220711 as a proxy for rs7209826 based on high LD (r^2^=0·99 in European ancestry populations), availability across most consortia, and prior functional evidence linking rs7220711 to *SOST* expression (see above). In addition, we validated the effect of rs7220711 on various measures of BMD as being comparable to that of rs7209826 (**Figure S3** in the appendix). There were no suitable proxies (r^2^>0·9) for rs188810925.

We examined the associations of rs7209826 and rs188810925 (and their selected proxies) on DXA-derived measures of BMD measured at specific body sites (lumbar spine, femoral neck, and forearm), using data from the largest GWAS to date for these phenotypes.^31^

We next examined the effect of these variants on *SOST* mRNA expression levels in various human tissues in the Genotype-Tissue Expression (GTEx) project dataset.^32^

### Study outcomes

Detailed definitions of each outcome studied are described in the **Supplementary Methods** and **Tables S3-5** in the appendix. We first tested the association of rs7209826 and rs188810925 with key efficacy outcomes, i.e., fracture risk and risk of osteoporosis (both defined as a combination of self-reported outcomes and International Classification of Diseases, ninth and tenth revision (ICD-9 and ICD-10) codes).

Next, we examined the association of the BMD-increasing alleles of rs7209826 and rs188810925 with risk of myocardial infarction and/or coronary revascularisation (including self-reported and ICD-9/ICD-10 codes for myocardial infarction, coronary artery bypass graft surgery and/or percutaneous transluminal coronary angioplasty) and a broader composite of all coronary heart disease (CHD; including all codes for myocardial infarction [MI] and/or coronary revascularisation, plus self-reported and ICD-9/ICD-10 codes for angina and chronic stable ischemic heart disease; see **Tables S3-5** in the appendix for specific codes included). Secondary disease outcomes included additional cardiovascular outcomes of interest (ischemic stroke, hemorrhagic stroke, all stroke, peripheral vascular disease, atrial fibrillation, heart failure), chronic kidney disease, aortic aneurysm and aortic stenosis (given sclerostin’s putative role in these conditions^17,33,34^), hypertension and T2D. Association with 11 quantitative cardiometabolic traits, many of which are established as causal risk factors for CHD, were also evaluated (systolic blood pressure (SBP), diastolic blood pressure (DBP), body mass index (BMI), waist-hip ratio (WHR) adjusted for BMI, low-density lipoprotein (LDL) cholesterol, high-density lipoprotein (HDL) cholesterol, triglycerides, fasting glucose, fasting insulin, HbA1c and serum creatinine-estimated glomerular filtration rate (eGFR)).

### Statistical Analyses

We derived cohort-specific SNP effect estimates from participants in the 4 prospective cohorts (see **Supplementary Methods** for specific methodology applied in each cohort). Estimates (log(OR) and the standard error of log(OR) for binary outcomes and beta and the standard error of beta for quantitative traits) for the per-allele effect of these variants on disease outcomes and quantitative traits were aligned to the BMD-increasing alleles (as per the effect of therapeutic sclerostin inhibition), and scaled to an increase in BMD equivalent to that reported with romosozumab treatment (see below). We then meta-analysed estimates from the prospective cohorts with (scaled) estimates from GWAS consortia for equivalent outcomes using inverse-variance weighted fixed-effect meta-analysis. We predefined a p-value threshold of <0·05 for the association with CHD given the reported association of sclerostin inhibition with cardiac ischemic events in prior RCTs.^13,14^ For the remaining 11 cardiometabolic outcomes and 11 quantitative traits, we set a Bonferroni adjusted p-value threshold of <0·0045 (0·05/11).

### Scaling of allelic estimates

We scaled allelic estimates pertaining to risk of osteoporosis, fractures, cardiometabolic outcomes and quantitative traits to an increase in BMD equivalent to that reported in a phase II RCT of 12 months of 210mg romosozumab monthly.^35^ This corresponds to the dose evaluated in phase III RCTs of romosozumab, and represents a 0·09 g/cm^2^ increase in lumbar spine BMD (LS-BMD) in postmenopausal women.^35^ See **Supplementary Methods** in the appendix for further details.

### Meta-analysis of cardiovascular outcomes in RCTs of sclerostin-inhibitors

We searched for all phase III RCTs performed for sclerostin inhibitors (see **Supplementary Methods** in the appendix for further details). We then performed meta-analyses of cardiovascular outcome data at 12 months from phase III RCTs using 210mg romosozumab monthly for 12 months, with ‘cardiac ischemic events’ as the primary outcome of interest. Further meta-analyses were performed for ‘cerebrovascular events’ and a composite outcome of ‘serious cardiovascular events’ (including cardiac ischemic events, cerebrovascular events, heart failure, cardiovascular death, non-coronary revascularisation and peripheral vascular ischemic events not requiring revascularisation). We set a prespecified p-value threshold of <0·05 for meta-analyses of the RCTs given the prior evidence for ischemic cardiovascular events seen in individual RCTs of romosozumab.^13,14^

All meta-analyses of RCT cardiovascular outcomes were performed according to the Mantel-Haenszel method without continuity correction,^36,37^ a non-parametric test designed for rare outcomes. Additional sensitivity analyses were performed using the Peto method^37^ (see **Supplementary Methods** in the appendix).

### Role of the funding source

The funders of the study had no role in study design, data collection, data analysis, data interpretation, or writing of the report. The corresponding authors (JB, CML and MVH) had full access to all the data in the study and shared final responsibility for the decision to submit for publication with all authors.

## Results

### Risk of cardiovascular events in phase III RCTs of sclerostin inhibitors

Romosozumab was the only sclerostin inhibitor with data from phase III RCTs. Four phase III RCTs of romosozumab, including 11,954 individuals were identified (**Table S6** in the appendix), of which three trials (BRIDGE^14^, ARCH^13^ and FRAME^12^) reported cardiovascular adverse event data (**Table S7** in the appendix). Only the BRIDGE^14^ and ARCH^13^ trials reported data on cardiac ischemic events and cerebrovascular events.

Meta-analysis of 25 cardiac ischemic events in 4,298 individuals, from two RCTs (BRIDGE^14^ and ARCH^13^) identified that 210mg romosozumab monthly, as compared to the comparator, led to a higher risk of disease (odds ratio [OR], 2·98; 95% confidence interval (CI), 1·18-7·55; P=0·02; **Figure 1**). Estimates from meta-analysis of ‘cerebrovascular events’ (27 events; OR, 2·15; 95% CI, 0·94-4·92; P=0·07) and ‘serious cardiovascular events’ (183 events; OR 1·21; 95% CI, 0·90-1·63; P=0·20) were directionally concordant with increased vascular risk arising from romosozumab treatment. Sensitivity analyses using the Peto-method showed similar results (**Figure S4** and **Table S8** in the appendix).

**Figure 1.**
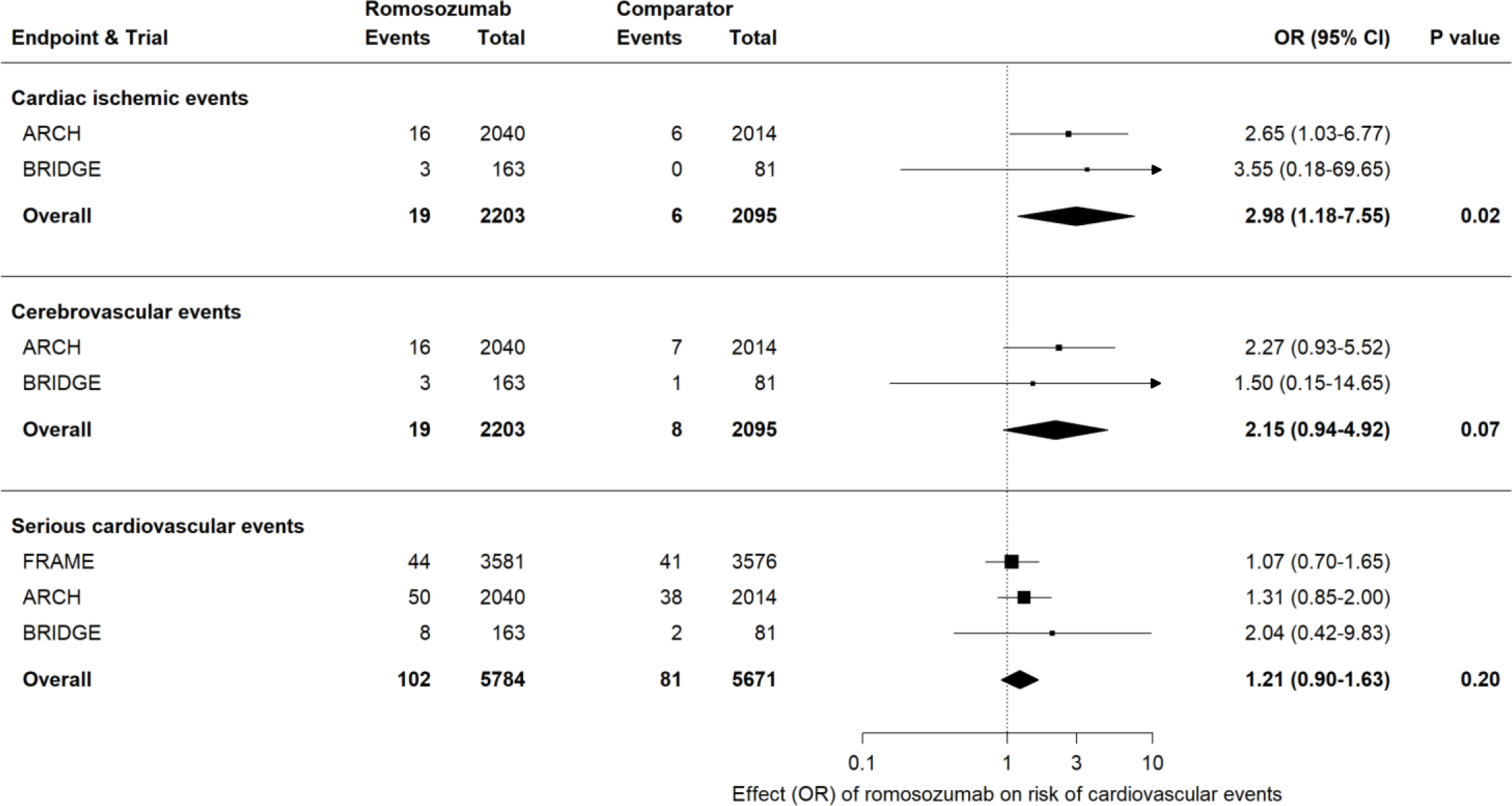
Meta-analysis of romosozumab and risk of cardiovascular events from phase III randomised controlled trials. *Events* represents number of events adjudicated per arm, during initial 12-month double-blind period in each trial. The *romosozumab*-group received 210mg romosozumab monthly in all trials; *comparator*-group received placebo (FRAME and BRIDGE trials) or alendronate (ARCH). Estimates derived using the Mantel-Haenszel method. Outcome data for *cardiac ischemic events* and *cerebrovascular events* were only available for the ARCH and BRIDGE trials. Boxes represent point estimates of effects. Lines represent 95% confidence intervals. OR, odds ratio; CI, confidence interval.

### Expression of *SOST* mRNA

Expression of *SOST* across 53 human tissue types in the Genotype-Tissue Expression consortium was highest in arterial tissue (**Figure S5** in the appendix; bone expression data not available). The minor alleles of rs7209826 (G-allele) and rs188810925 (A-allele) were both associated with lower expression of *SOST* mRNA in various human tissues, with the strongest associations with *SOST* expression for each variant identified in tibial artery (rs7209826: P=1·4×10^−8^) and aorta (rs188810925; P=7·6×10^−6^; **Figure S6** in the appendix).

### Association of rs7209826 and rs188810925 with BMD

The minor alleles of both SNPs were associated with higher estimated heel-bone BMD (eBMD) in UKBB (rs7209826: 0·04 g/cm^2^ [95% CI, 0·04-0·05; P=2·3×10^−36^] per G allele and rs188810925: 0·07 g/cm^2^ [95% CI, 0·05-0·08; P=1·3×10^−26^] per A allele, **Figure S7** in the appendix), and with higher LS-BMD (rs7209826: 0·008 g/cm^2^ [95% CI, 0·005-0·01; P=5·4×10^−07^] per G allele; rs188810925: 0·016 g/cm^2^ [95% CI, 0·01-0·022; P=4·3×10^−07^] per A allele, **Figure S7** in the appendix). We subsequently scaled all further genetic estimates to the increase in lumbar spine BMD seen with 12 months of 210mg romosozumab monthly (equivalent to a 0·09 g/cm^2^ higher BMD).^35^

### Association with osteoporosis and fracture risk

Scaled to match the effect of 12 months of 210mg romosozumab monthly on lumbar spine BMD, meta-analysis of rs7209826 and rs188810925 yielded a 57% lower risk of osteoporosis (OR, 0·43; 95% CI, 0·36-0·52; P=2·4×10^−18^) and a 41% lower risk of sustaining a bone fracture (OR, 0·59; 95% CI, 0·54-0·66; P=1·4×10^−24^; **Figure 2** and **Figure S8** in the appendix). Associations were consistent across fracture sites (**Figure S9** in the appendix).

**Figure 2.**
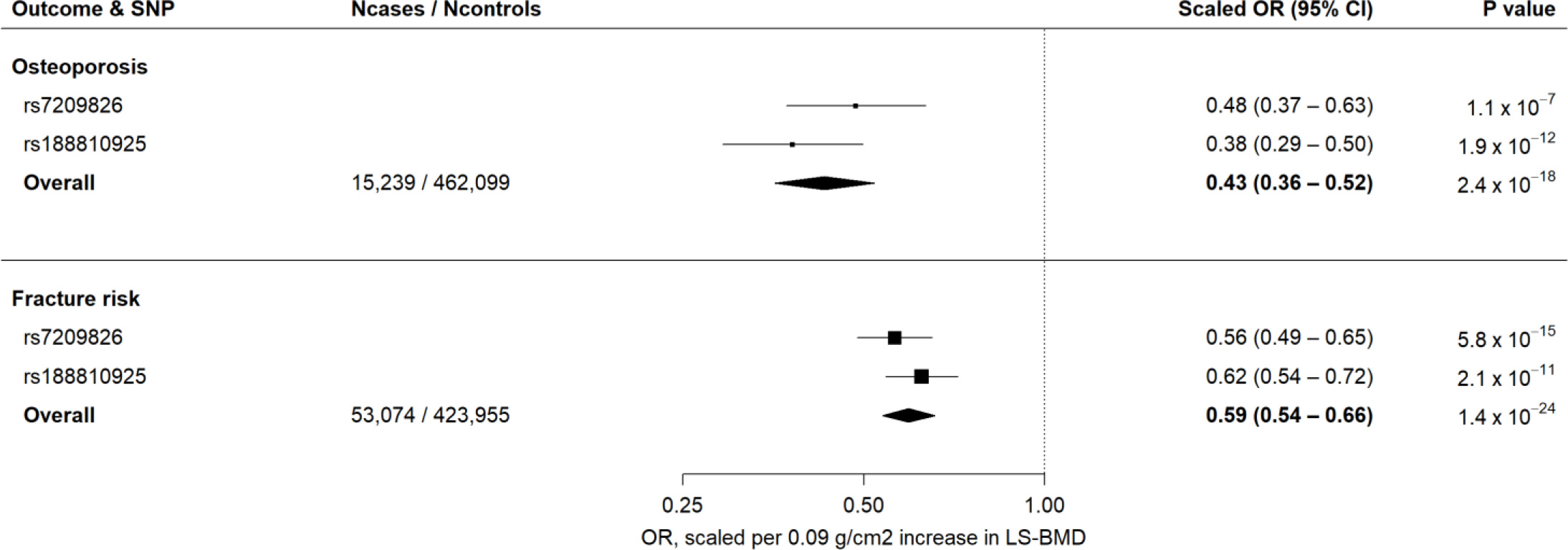
Scaled estimates and meta-analysis of BMD-increasing SOST variants with risk of osteoporosis (15,329 cases) and fracture (53,074 cases). Estimates are scaled to match the effect of 210mg romosozumab monthly for 12 months on lumbar spine bone mineral density (0·09 g/cm^2^; see Methods) and aligned to the BMD-increasing alleles. See Methods sections for outcome definitions. Boxes represent point estimates of effects. Lines represent 95% confidence intervals. OR, odds ratio; CI, confidence interval; LS-BMD, lumbar spine bone mineral density.

### Association with coronary heart disease

In meta-analysis of scaled estimates including up to 69,649 cases, the BMD-increasing *SOST* variants associated with an 18% higher risk of myocardial infarction and/or coronary revascularisation (OR, 1·18; 95% CI, 1·06-1·32; P=0·003; **Figure 3** and **Figure S10** in the appendix). Using a broader definition of CHD, which included self-reported angina and chronic stable ischemic heart disease (up to 106,329 cases), the *SOST* variants associated with a 13% increased risk of disease (OR, 1·10; 95% CI, 1·00-1·20; P=0·04; **Figure 3** and **Figure S10** in the appendix).

**Figure 3.**
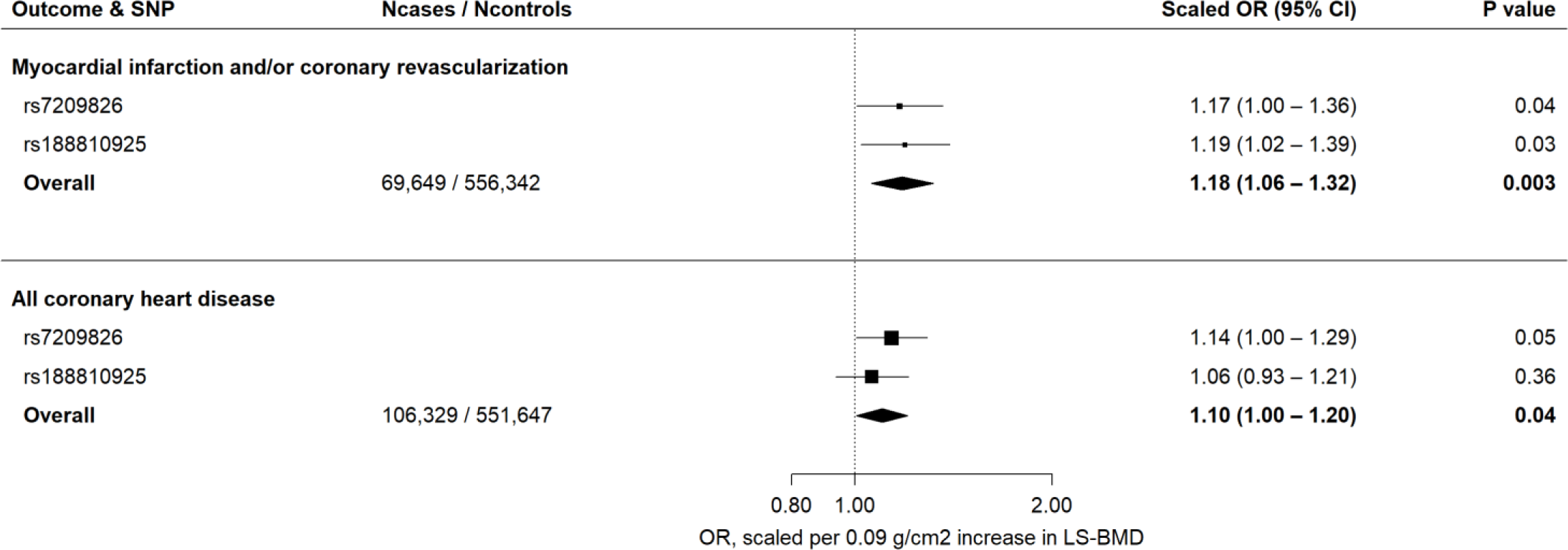
Scaled estimates and meta-analysis of BMD-increasing *SOST* variants with risk of myocardial infarction and/or coronary revascularisation (up to 69,649 cases) and coronary heart disease (up to 106,329 cases). Estimates are scaled to match the effect of 210mg romosozumab monthly for 12 months on lumbar spine bone mineral density (0·09 g/cm^2^; see Methods) and aligned to the BMD-increasing alleles. Boxes represent point estimates of effects. Lines represent 95% confidence intervals. OR, odds ratio; CI, confidence interval.

### Association with additional cardiometabolic risk factors and diseases

Evaluation of cardiometabolic risk factors and diseases revealed that the BMD-increasing *SOST* variants associated with higher risks of hypertension (OR, 1·12; 95% CI, 1·05-1·20; P=8·9×10^−4^) and T2D (OR, 1·15; 95% CI, 1·05-1·27; P=0·003; **Figure 4A**).

**Figure 4.**
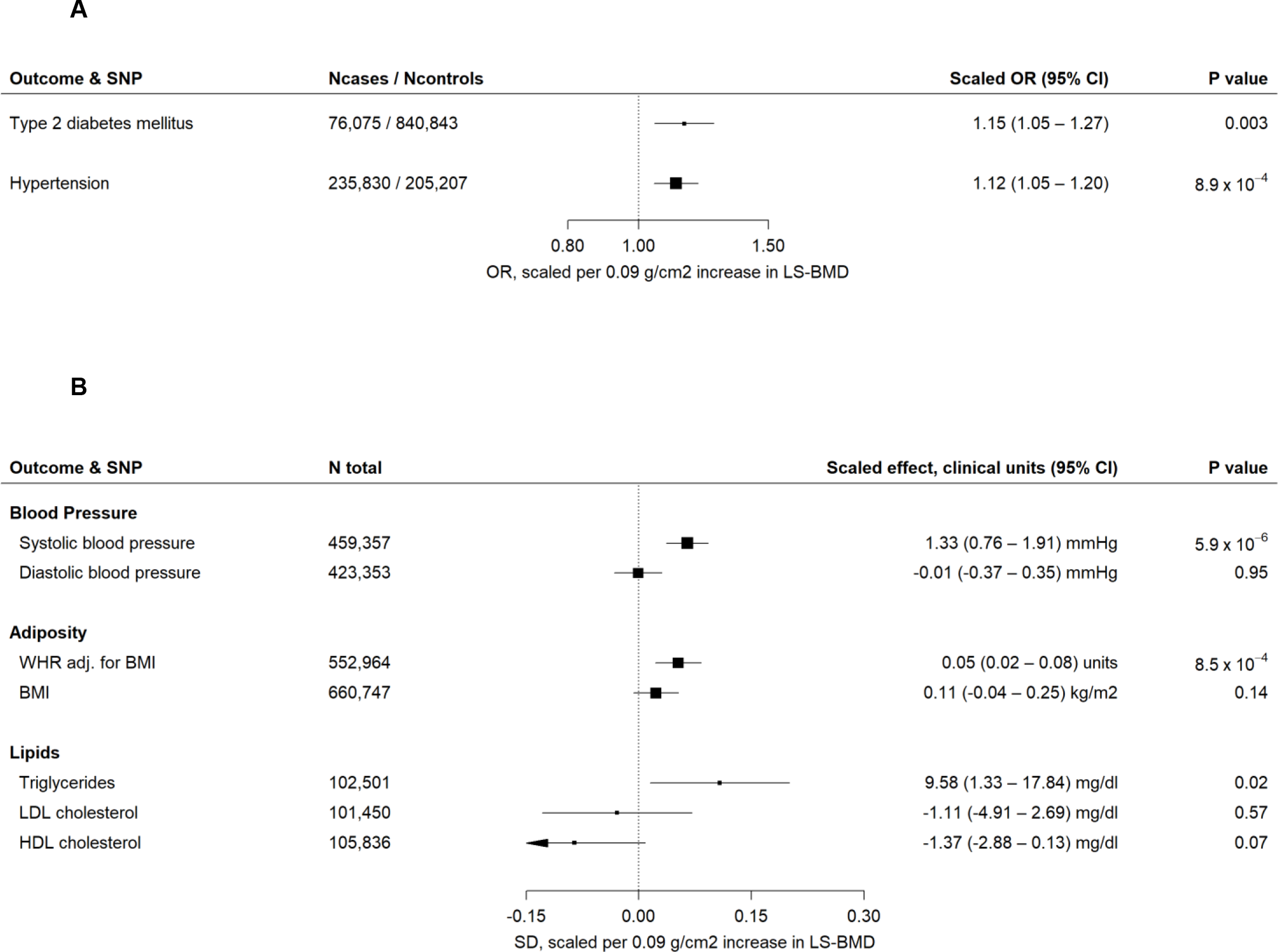
Meta-analysis of BMD-increasing *SOST* variants and cardiometabolic risk factors. Panel A shows association with hypertension and type 2 diabetes mellitus; Panel B shows association with quantitative traits plotted in SD units, with clinical units for each trait indicated in the column on the right. All estimates are scaled to match the effect of 210mg romosozumab monthly for 12 months on lumbar spine bone mineral density (0·09 g/cm^2^; see Methods) and aligned to the BMD-increasing alleles. The significance threshold was set at 0·0045 (see Methods for details). Boxes represent point estimates of effects in odds ratio (panel A) or standard deviation (panel B) units. Lines represent 95% confidence intervals. UKBB, UK Biobank; OR, odds ratio; CI, confidence interval; mmHg, millimetres of mercury; SD, standard deviations, WHR, waist to hip ratio; adj, adjusted; BMI, body mass index; LDL, low-density lipoprotein; HDL, high-density lipoprotein.

Consistent with the effect on hypertension, we found the *SOST* variants to be associated with 1·3 mmHg higher SBP (95% CI, 0·8-1·9; P=5·9×10^−6^), but observed no effect on DBP (**Figure 4B**). In addition, the *SOST* variants associated with 0·05 standard deviation (SD) units higher WHR adjusted for BMI (95% CI, 0·02-0·08; P=8·5×10^−4^), and, nominally, with higher serum triglycerides (9·6 mg/dl higher; 95% CI, 1·3-18·8 mg/dl; P=0·02). We found no further associations (**Figures S11-14** and **Tables S9-10** in the appendix).

### Trans-ethnic replication

We attempted trans-ethnic replication in the China Kadoorie Biobank. In 21,547 individuals with eBMD measurements available, the rs7209826 variant showed no evidence of association (beta=0·0016 g/cm^2^; SE=0·001; P=0·14) and strong evidence of heterogeneity with the corresponding eBMD estimate in UKBB (P-heterogeneity=1·35×10^−28^; **Figure S15** in the appendix). A regional analysis showed no evidence of a suitable genetic instrument (**Figure S16** in the appendix). Of note, the lack of association with CHD and other endpoints (**Table S11** in the appendix) in CKB suggests that the associations obtained in Europeans were not due to pleiotropy.

### Triangulation of randomised controlled trials and human genetics

We performed fixed-effect meta-analysis of 337 clinical fractures in 11,273 individuals from two RCTs with fracture outcome data (FRAME^12^ and ARCH^13^, **Table S12** in the appendix), and identified that 210mg romosozumab monthly, as compared to comparator, led to a 32% lower risk of clinical fracture (hazard ratio 0·68; 95% CI, 0·55-0·85; P=6·0×10^−4^; **Figure 5** and **Figure S17** in the appendix) at 12 months. This compared well to the scaled genetic estimate, using the two *SOST* variants, of a 41% lower risk of fracture (OR, 0·59, 95% CI, 0·54-0·66, P= 1·4×10^−24^). Evidence from treatment trials (two romosozumab phase III RCTs^13,14^), and human genetics (two *SOST* variants) showed that inhibition of sclerostin increases risk of coronary events **(Figure 5)**, indicating that this adverse effect is likely real and target-mediated.

**Figure 5.**
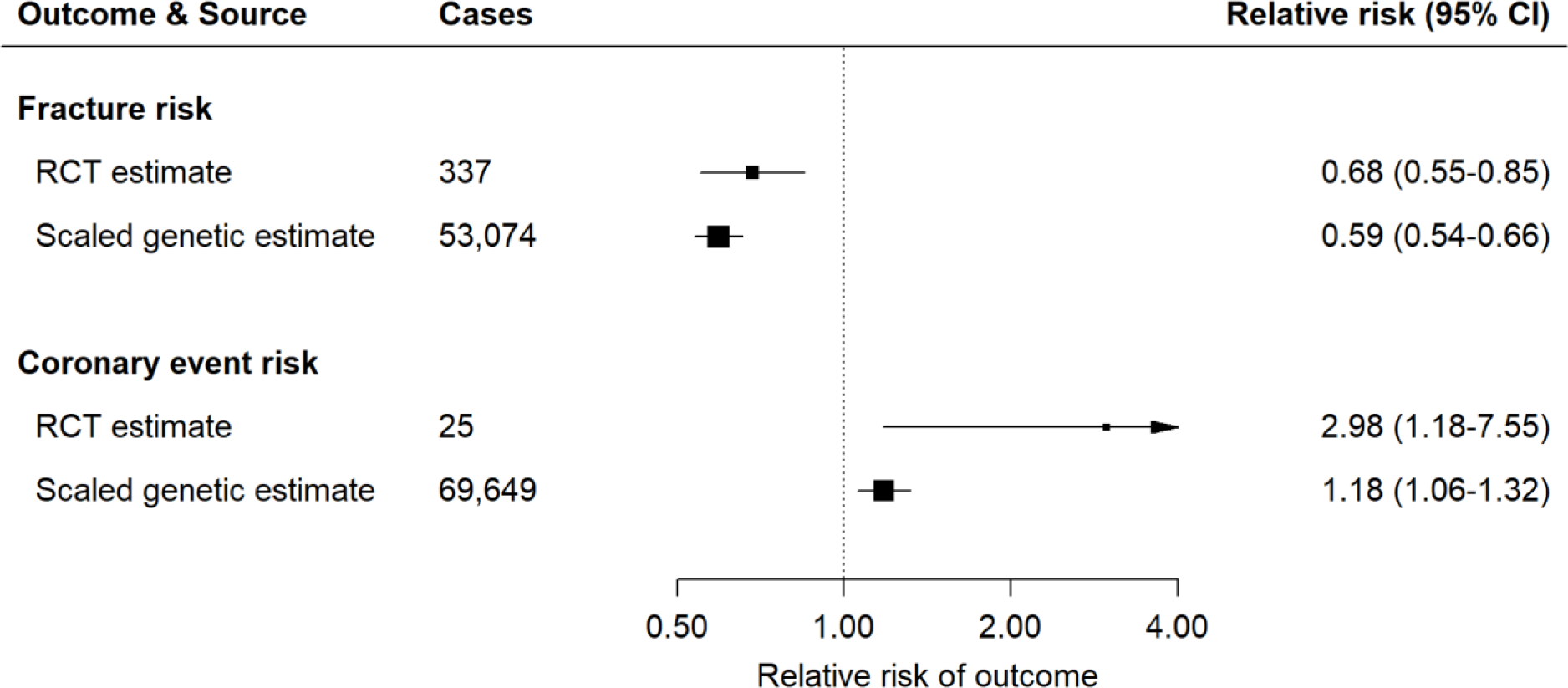
Inhibition of sclerostin and risk of fracture and coronary events derived from meta-analysis of phase III randomised controlled trials of romosozumab and human genetics. *Fracture risk RCT estimate* represents inverse variance weighted fixed-effect meta-analysis of estimates for hazard ratio of ‘clinical fracture’ at 12 months (a composite of non-vertebral or symptomatic vertebral fracture) in the ARCH and FRAME trials. *Coronary event risk RCT* estimate represents fixed-effect meta-analysis (using the Mantel-Haenszel method) of estimates for odds ratio of ‘cardiac ischemic events’ in the ARCH and BRIDGE trials. *Scaled genetic estimates for fracture risk* and *coronary event risk* represent inverse variance weighted fixed-effect meta-analyses of the scaled allelic estimates for the odds ratio of fracture risk and myocardial infarction and/or coronary revascularisation, respectively. All RCT estimates refer to the effect of romosozumab 210mg monthly for 12 months relative to comparator, and all genetic estimates are scaled to match the effect of 210mg romosozumab monthly for 12 months on lumbar spine bone mineral density (0·09 g/cm^2^ increase). Boxes represent point estimates of effects. Lines represent 95% confidence intervals. RR, relative risk; CI, confidence interval.

## Discussion

Using data from both naturally-occurring human genetics and randomised interventional trials, we have shown that while raising BMD through lifelong lowering of sclerostin is associated with lower risk of osteoporosis and fractures, it also results in a higher risk of CHD. While prior phase III RCTs of sclerostin inhibition by romosozumab suggested an increased risk of adverse cardiac events, it remained possible that the finding was due to chance owing to the low number of events (only 25 cases). Our genetic analysis, including up to 106,329 CHD cases, shows that this excess risk of CHD from sclerostin inhibition is very likely to be real. Our findings also suggest that the excess risk of coronary events may be driven by an increase in central adiposity, SBP (and hypertension), and increased risk of T2D.

There are several additional lines of evidence supporting the findings of our study. Sclerostin is expressed in cardiovascular tissues,^33,38^ supporting a potential biological role in these tissue types. In observational studies, higher levels of circulating sclerostin are associated with a higher risk of cardiovascular disease together with higher levels of cardiometabolic risk factors such as hypertension, T2D and central adiposity.^39–44^ However, further evidence has suggested that sclerostin may be up-regulated in the vasculature in response to vascular calcification, as part of a regulatory process aimed at counteracting such calcification.^17^ Recent murine studies have implicated sclerostin in adipocyte metabolism^45^ and the development of atherosclerosis, aortic aneurysm and hemopericardium.^34,46^ However, extrapolating findings from animal models of disease to humans is plagued by failures of translation^47,48^, and observational studies of humans can be influenced by sources of error: this is why studies employing a randomised design in humans provide more reliable evidence on causation.^49^

Elucidation of whether adverse effects are on- or off-target is critical^8^ as on-target effects would mean that any drug under development in the same class (i.e., a sclerostin inhibitor), would be expected to share a similar adverse effect profile (i.e. higher risk of vascular events). Our genetic data provide strong evidence in support of target-mediated adverse effects of sclerostin inhibition on coronary events. Sclerostin exerts its effects on bone as an inhibitor of the Wnt-signaling pathway (a pathway also previously linked to vascular calcification^50–52^) by binding to the Wnt co-receptor low density lipoprotein receptor-related protein (LRP) 5 and 6.^53^ Protein-coding mutations in both *LRP5* and *LRP6* have been linked to alterations in BMD and cardiometabolic risk profiles, including insulin resistance, dyslipidemia, hypertension and CHD,^54,55^ which supports our findings of altered cardiometabolic disease in carriers of *SOST* variants.

Concerns of cardiovascular safety have also plagued other anti-osteoporotic agents. Odanacatib, a cathepsin K inhibitor developed by Merck for the treatment of osteoporosis, while shown to reduce fracture risk in phase III trials, was not developed further owing to an increased risk of stroke.^56^ While we did not identify strong associations of *SOST* with risk of stroke subtypes in our analyses, the point estimates for both ischemic and hemorrhagic stroke were OR > 1. With genetic lowering of sclerostin leading to elevations in systolic blood pressure and WHR adjusted for BMI (each of which plays a causal role in stroke^57,58^), it is plausible that additional stroke cases would identify such an association. More broadly, genetically elevated BMD may exert a modest causal effect on risk of T2D and CHD.^59^ Interestingly, other BMD-raising therapies, e.g., denosumab or bisphosphonates, have not shown an association with increased cardiovascular risk in clinical trials.^60–62^

The approach to exploiting human genetics to validate target-mediated effects is well-established.^22–26^ In our study, we selected variants associated with reduced expression of *SOST* and increased BMD as proxies for the effect of sclerostin inhibition. We validated the effects of the *SOST* genetic variants on risk of osteoporosis and bone fracture (including fracture across several sites), and leveraged a large number of CHD cases (a more than 2,700-fold increase over the number of cases reported in phase III trials of romosozumab^13,14^). Our approach also facilitated identification of potential mechanisms and mediators that may drive this excess coronary risk. For example, the association of *SOST* variants with central adiposity, SBP and T2D has not been previously reported and if not measured in a clinical trial, such associations cannot be readily explored. Whilst trans-ethnic replication in a large East Asian biobank (CKB) could not be reliably performed owing to a lack of association of *SOST* variants with BMD, we note the lack of association with CHD and other outcomes in CKB (**Table S11** in the appendix). This provides an opportune means to illustrate that the multiple cardiometabolic risk factor and disease associations identified in the European datasets, where genetic variants in *SOST* do act as valid instruments for inhibition of sclerostin, did not arise as a consequence of horizontal pleiotropy (in other words, the genetic associations did not arise through a mechanism other than that which operates through inhibition of sclerostin).^63^

Our findings warrant a rigorous assessment of the effect of romosozumab (and other sclerostin inhibitors in clinical development) on cardiovascular disease and cardiometabolic risk factors, particularly in light of the ongoing licensing applications for this drug. Given the high short-term mortality associated with some types of fragility fracture (e.g., up to 25% mortality in the 12 months following a hip fracture^64^), a risk-benefit assessment is warranted, weighing the merits of lower fracture (and fracture-related morbidity and mortality) against the potential harm from higher risk of metabolic and vascular disease.

In conclusion, we have shown that genetic variants in the *SOST* locus that mimic pharmacological inhibition of sclerostin are associated with lower *SOST* expression, lower risk of osteoporosis and fractures but an elevated risk of coronary heart disease, likely explained by increases in systolic blood pressure, central adiposity and risk of type 2 diabetes mellitus. This adds valuable information as to whether pharmacological inhibition of sclerostin should be pursued as a therapeutic strategy for the prevention of fracture.

## Declaration of Interests

BMN is a member of the Scientific Advisory Board for Deep Genomics, a consultant for Camp4 Therapeutics Corporation, a consultant for Merck & Co., a consultant for Takeda Pharmaceutical, and a consultant for Avanir Pharmaceuticals, Inc. MVH has collaborated with Boehringer Ingelheim in research, and in accordance with the policy of the Clinical Trial Service Unit and Epidemiological Studies Unit (University of Oxford), did not accept any personal payment. All other authors declare no competing interests.

## Acknowledgements

JB is supported by funding from the Rhodes Trust, Clarendon Fund and the Medical Sciences Doctoral Training Centre, University of Oxford. JCC is funded by the Oxford Medical Research Council Doctoral Training Partnership (Oxford MRC DTP) and the Nuffield Department of Clinical Medicine, University of Oxford. TF is supported by the NIHR Biomedical Research Centre, Oxford. SLP has a Veni Fellowship (016.186.071; ZonMW) from the Dutch Organization for Scientific Research, Nederlandse Organisatie voor Wetenschappelijk Onderzoek (NWO). LM Is supported by funding from Estonian Research Council Grants PRG184, IUT20-60, IUT34-4, IUT34-11, IUT24-6. B.M.N. is supported by NIH project 5P50HD028138-27-Project 2: Common Complex Trait Genetics of Reproductive Phenotypes. RM Is supported by funding from Estonian Research Council Grants PRG184, IUT20-60, IUT34-4, IUT34-11, IUT24-6. CML is supported by the Li Ka Shing Foundation; WT-SSI/John Fell funds; the NIHR Biomedical Research Centre, Oxford; Widenlife; and NIH (5P50HD028138-27). MVH works in a unit that receives funding from the UK Medical Research Council and is supported by a British Heart Foundation Intermediate Clinical Research Fellowship (FS/18/23/33512) and the National Institute for Health Research Oxford Biomedical Research Centre. China Kadoorie Biobank (CKB) is supported by funding from Kadoorie Charitable Foundation in Hong Kong; the UK Wellcome Trust (grant numbers 202922/Z/16/Z, 088158/Z/09/Z, 104085/Z/14/Z); Chinese Ministry of Science and Technology (grant number 2011BAI09B01); National Natural Science Foundation of China (Grant numbers 81390540, 81390541, 81390544); UK Medical Research Council (grant numbers MC_PC_13049, MC_PC_14135); GlaxoSmithKline; BHF Centre of Research Excellence, Oxford (grant number RE/13/1/30181); British Heart Foundation; and Cancer Research UK. CKB wishes to thank the CKB participants; CKB project staff based at Beijing, Oxford and the 10 regional centres; the China National Centre for Disease Control and Prevention (CDC) and its regional offices for assisting with the fieldwork, and the assistance of BGI (Shenzhen, China) for conducting DNA extraction and genotyping.

