## Supplemental Appendix for "Lifelong genetically lowered sclerostin and risk of cardiovascular disease"


**Supplementary Methods** 2

Detailed cohort information 2

UK Biobank (UKBB) 2

Partners HealthCare Biobank (PHB) 4

Estonian Biobank (EGCUT) 6

China Kadoorie Biobank (CKB) 6

Genome-wide association study (GWAS) consortia 8

Selection of proxy SNPs 8

Scaling of allelic estimates 8

Meta-analysis of scaled allelic estimates 8

Identification of phase III clinical trials of sclerostin inhibitors 8

Meta-analysis of randomized controlled trials 8

**Supplementary Figures** 9

Figure S1. Overview of study design. 9

Figure S2. Regional association plots of SOST locus in GWAS of heel-bone estimated bone mineral density in UKBB (n = 142,487). 10

Figure S3. Per-allele associations of rs7209826 and selected proxy (rs7220711; r2 = 0.99) with various bone mineral density (BMD) measures. 11

Figure S4. Meta-analysis of romosozumab and risk of cardiovascular events from phase III randomized controlled trials, using Peto Method. 12

Figure S5. SOST expression is highest in arterial tissues. 13

Figure S6. SOST mRNA expression by rs188810925 and rs7209826 genotype. 14

Figure S7. Per-allele associations of rs7209826 and rs188810925 with various bone mineral density (BMD) measures. 15

Figure S8. Study-specific scaled estimates and meta-analysis of BMD-increasing SOST variants with risk of osteoporosis and fracture. 16

Figure S9. Scaled estimates and meta-analysis of BMD-increasing SOST variants with risk of fracture in the preceding 5 years, at different anatomical sites, in UKBB. 17

Figure S10. Study-specific scaled estimates and meta-analysis of BMD-increasing SOST variants with risk of myocardial infarction and/or coronary revascularization and coronary heart disease. 18

Figure S11. Study-specific scaled estimates and meta-analysis of BMD-increasing SOST variants with risk of 11 cardiometabolic outcomes. 19

Figure S12. Study-specific scaled estimates and meta-analysis of BMD-increasing SOST variants with systolic and diastolic blood pressure. 20

Figure S13. Study-specific scaled estimates and meta-analysis of BMD-increasing SOST variants with waist to hip ratio (WHR; adjusted for body mass index), body mass index (BMI), LDL cholesterol, HDL cholesterol and triglycerides. 21

Figure S14. Scaled estimates and meta-analysis of BMD-increasing SOST variants with glomerular filtration rate estimated from serum creatinine (eGFR) in the CKDGEN consortium. 22

Figure S15. Per-allele association of rs7209826 with estimated heel-bone bone mineral density (eBMD) in UKBB and China Kadoorie Biobank. 23

Figure S16. Regional association plot of SOST locus in GWAS of heel-bone estimated bone mineral density in China Kadoorie Biobank (n = 21,547). 24

Figure S17. Meta-analysis of romosozumab and risk of clinical fracture from phase III randomized controlled trials. 25

**Supplementary Tables** 26

Table S1. Baseline characteristics in UK Biobank. 26

Table S2. Genome-wide association studies and consortia included in present study. 27

Table S3. Definitions of outcomes analysed in UK Biobank. 28

Table S4. Definitions of outcomes analysed in Partners HealthCare Biobank. 29

Table S5. Definitions of outcomes analysed in China Kadoorie Biobank. 30

Table S6. Published phase III randomized controlled trials of romosozumab. 31

Table S7. Reported cardiovascular outcomes at 12 months from published phase III randomized controlled trials of romosozumab. 32

Table S8. Summary statistics of meta-analyses of cardiovascular events in randomized controlled trials of romosozumab. 33

Table S9. Scaled estimates for glycaemic traits from MAGIC consortium. 34

Table S10. Summary statistics of meta-analyses of scaled allelic estimates. 35

Table S11. Per-allele estimates for rs7209826 in China Kadoorie Biobank. 36

Table S12. Reported fracture outcomes at 12 and 24 months from published phase III randomized controlled trials of romosozumab. 37

**Supplementary References** 37

### Supplementary Methods

#### Detailed cohort information

##### UK Biobank (UKBB)

##### Study population in UKBB

UK Biobank (UKBB) is a prospective study of more than 500,000 British individuals recruited from 2006 to 2010 and aged between 45 and 69.[1] Phenotypic information available includes self-reported medical history as ascertained by verbal interview with a medical professional at enrolment, hospital-derived electronic health record (EHR) data, including International Classification of Diseases, ninth and tenth revision (ICD-9 and ICD-10) codes and Office of Population and Censuses Surveys (OPCS-4) procedure codes, and an extensive set of physical measurements. Genotypic data (see below for detail) is available for 488,377 individuals, of whom 487,409 have imputed genotype data.

For the purposes of this study, we excluded all samples indicated to have poor quality genotypes by UKBB (on the basis of high sample heterozygosity and missingness), and further excluded individuals that had: reported non-white or mixed ethnicity at any point during follow-up; withdrawn their consent for participation; >10 third degree relatives; putative sex chromosome aneuploidy; sex mismatches (comparing genetically determined vs. self-reported sex, and comparing between assessments); ethnicity mismatches (mismatches between genetically determined and self-reported ethnicity for white British individuals, and any ethnicity mismatches between assessments). We reviewed pairwise genetic relatedness between individuals and excluded one individual per pair of individuals with an estimated 2nd degree or closer relatedness (equivalent to a kinship coefficient of greater than 0.0884[2]). After applying these filters, 423,761 subjects remained. Baseline characteristics of participants in UKBB are shown in Table S1.

##### Genotyping in UKBB

Genotyping, quality control and imputation were performed centrally by UKBB, and details are fully described elsewhere.[1] Briefly, genotype data is available for 488,377 individuals, 49,950 of whom were genotyped using the Applied Biosystems™ UK BiLEVE Axiom™ Array by Affymetrix (containing 807,411 markers[3]). The remaining 438,427 individuals were genotyped using the Applied Biosystems™ UK Biobank Axiom™ Array by Affymetrix (containing 825,927 markers). Both of these arrays were specifically designed for use in the UKBB project and share ~95% of marker content. Phasing was done using SHAPEIT3, and imputation was conducted using IMPUTE4. For imputation, the Haplotype Reference Consortium (HRC) panel[4] was used wherever possible, and for single nucleotide polymorphisms (SNPs) not in that reference panel, a merged UK10K + 1000 Genomes reference panel was used. SNPs were imputed from both panels, but the HRC imputation was preferentially used for SNPs present in both panels.

Both SNPs selected as instruments in this study demonstrated high imputation quality in UKBB (info scores of 0.98 and 0.95 for rs7209826 and rs188810925, respectively).

##### Study outcomes and definitions in UKBB

*Fracture:* Fracture status was based on an affirmative answer to the question "Have you fractured/broken any bones in the last 5 years?" at either baseline or at follow-up. Individuals were designated as missing if they answered "Do not know" or "Prefer not to answer" at baseline and at all follow-up occasions. All remaining individuals were designated as controls. An identical definition has been used in recent GWAS of BMD and fracture,[5,6] and the use of self-reported fracture has previously been validated.[7] Individuals who reported having sustained a fracture were asked "Which bone(s) did you fracture/break?", and given a list of option to choose from (including ankle, leg, hip, spine, wrist, arm and other bones). For the purposes of our analysis, we grouped these into upper limb (wrist, arm), lower limb (ankle, leg, hip), vertebral (spine) and other fractures.

*Bone mineral density*: Detailed descriptions pertaining to measurement of eBMD and BMD are provided elsewhere.[6,8] Estimates for eBMD were provided in g/cm$^2$ units.[6] Estimates for DXA-derived BMD across three anatomical sites (lumbar spine (LS-BMD), femoral neck (FN-BMD) and forearm (FA-BMD)) were provided in standard deviation (SD) units.[8] We converted these estimates to g/cm$^2$ units using an estimate of the pooled population SD for each these three measures (0.18 g/cm$^2$, 0.14 g/cm$^2$ and 0.07 g/cm$^2$ for LS-BMD, FN-BMD and FA-BMD, respectively) in the GEFOS consortium.[8]

*Systolic and diastolic blood pressure*: Individual values for systolic and diastolic blood pressure and the definition of hypertension were derived using the same method as in a recent GWAS for blood pressure performed in UKBB.[9] We derived the mean SBP and DBP for each subject from two blood pressure measurements performed at baseline. For subjects with only a single measurement available, we used this single value. We then adjusted these values for blood pressure-lowering medication use by addition of 10 and 15 mmHg to DBP and SBP, respectively, for subjects who had reported taking blood pressure medication at baseline.[10] Individuals were coded as having hypertension if they had an adjusted SBP of 140 mmHg or higher, DBP of 90 mmHg or higher, or reported use of antihypertensive medication.

*Adiposity*: To create phenotypes for body mass index (BMI) and waist-to-hip ratio (WHR) adjusted for BMI (WHRadjBMI), we followed steps consistent with those followed in the GIANT consortium.[11–13] Baselines measures of BMI, waist circumference and hip circumference were used. We first divided waist circumference by hip circumference to calculate the WHR, and then regressed WHR on BMI, sex, age at time of assessment and (age at time of assessment)$^2$, whereas BMI was regressed on sex, age at time of assessment and (age at time of assessment)$^2$. Next we performed rank inverse normalization on the resulting residuals from the regressions. These normalized residuals (for BMI and WHRadjBMI) were used for performing allelic association testing.

*Coronary heart disease*: We included two definitions for CHD: an inclusive CHD phenotype including angina and other forms of chronic CHD, and a more specific phenotype of myocardial infarction and/or coronary revascularization only. Both of these definitions were identical to those used in a recent GWAS of CHD conducted in UKBB by the CARDIoGRAMplusC4D consortium.[14]

Detailed definitions for all binary outcomes analysed in UKBB (and not specifically defined above) are given in Table S3.

**Association analyses in UKBB**

The genotype-outcome association analyses in UKBB were performed using SNPTEST v2.5.4. We used an additive frequentist model (using "-*frequentist 1*") and included sex, age at baseline, genotyping array (a binary variable), and the first 15 principal components as covariates in all analyses. We accounted for genotype uncertainty by using *"-method expected"*.

**Ethical considerations in UKBB**

The UKBB project was approved by the North West Multi-Centre Research Ethics Committee and all participants provided written informed consent to participate. This research has been conducted under UKBB application number 11867.

**Partners HealthCare Biobank (PHB)**

**Study population in PHB**

We identified patients with relevant phenotype information and genotype data from the Partners HealthCare Biobank,[15,16] a biorepository of consented patient samples at Partners HealthCare hospitals in the Boston area of Massachusetts. The Partners HealthCare Biobank maintains blood and DNA samples, clinical records and genotype data from consented patients. Patients are recruited in the context of clinical care appointments and electronically. All patients who participate in the Partners Biobank are consented for their samples to be linked to their clinical information for the use in broad-based research.

A total of 19,136 patients of European ancestry were genome-wide genotyped in Partners Biobank, including 8,868 males and 10,268 females with age at recruitment ranged from 19 to 102 (mean=59.4, median=62, SD=16.61). For these patients, electronic health records were available for extracting the following phenotypes: fracture, osteoporosis, coronary heart disease, myocardial infarction and/or coronary revascularization, type 2 diabetes, body mass index, HDL-cholesterol, LDL-cholesterol, triglycerides, and hypertension. We subset these patients according to the phenotype information availability for each genetic association analysis (See *Study outcomes and definitions in PHB* below).

**Genotyping population in PHB**

DNA samples from whole blood and genome-wide genotyping were done for 25,582 patients. Genotyping were done with Illumina Multi-Ethnic Genotyping Array (first batch), Expanded Multi-Ethnic Genotyping Array (second batch), and Multi-Ethnic Global BeadChip (third batch), all of which were designed to capture the global diversity of genetic backgrounds (N of genotyped variants: 1,416,020 – 1,778,953).

Pre-imputation QC was performed on each genotyping batch separately as follows: we removed single nucleotide polymorphisms (SNPs) with genotype missing rate > 0.05 before sample-based QC; excluded samples with genotype missing rate > 0.02, absolute value of heterozygosity > 0.2, or failed sex checks; removed SNPs with missing rate > 0.02 after sample-based QC. To merge genotyping batches for imputation and analyses, we performed batch QC by removing SNPs with significant batch association (p-value < $1.0\text{x}10^{-6}$ between different batches).

Since the Partners Biobank samples have diverse population backgrounds, we performed Hardy-Weinberg equilibrium test (p-value < $1.0\text{x}10^{-6}$) for SNP-based QC after extracting samples of European ancestry (see below). We also performed relatedness tests by identifying pairs of samples with $\pi > 0.2$ and excluding one sample from each related sample pair (560 samples excluded). All QC were conducted using PLINK v1.9 and R software.

We extracted samples with European ancestry based on principal component analysis (PCA) with 1000 Genomes Project reference samples. To identify patients of European ancestry, we first performed PCA on LD-pruned dataset merged with 1000 Genomes Project reference samples labeled with 5 distinct super-populations (European [EUR], African [AFR], East Asian [EAS], admixed American [AMR], and South Asian [SAS]). Then, we used the top 4 PCs to build a random forest classifier trained on 1000 Genomes Project reference samples with super-population labels. Finally, we applied the trained random forest classifier to identify patients of European ancestry from Partners Biobank (with predicted probability of European ancestry > 0.9).

Genotype imputation was performed on the QCed patients of European ancestry with a 2-step pre-phasing/imputation approach. We used Eagle2 for the pre-phasing and minimac3 for imputation, with a reference panel from 1000 Genomes Project phase 3.

**Study outcomes and definitions in PHB**

We extracted relevant phenotype information from electronic health records for the patients with imputed genome-wide genotype data. The phenotype definition and number of samples are described in Table S4.

**Association analyses in PHB**

Linear (for continuous phenotype) or logistic regression (for case-control binary phenotype) was used to test associations between SNPs rs188810925 and rs7209826 and phenotypes of interest using PLINK 1.9. For phenotypes without residual inverse rank normalization, we adjusted for sex, age at assessment, age squared, genotype batches and 15 principal components in the regression model. For phenotypes with residual inverse rank normalization, we did not adjust for any covariates in the regression model.

**Ethical considerations in PHB**

The Partners HealthCare Biobank maintains blood and DNA samples from consented patients seen at Partners HealthCare hospitals in the Boston area of Massachusetts. Patients are recruited in the context of clinical care appointments, and also electronically. Biobank subjects provide consent for the use of their samples and data in broad-based research.

**Estonian Biobank (EGCUT)**

**Study population in EGCUT**

The Estonian Biobank is the population-based biobank containing longitudinal data and biological samples, including DNA, for 5% of the adult population of Estonia. The broad informed consent form signed by the participants of the biobank allows the Estonian Genome Center to continuously update their records through periodical linking to central electronic health record databases and registries. We studied the genotypic and phenotypic data of 51881 individuals and after removing relatives (PiHat >0.2), 36073 individuals (65% women, 35% men) with average age of 45 years were included for further analysis.

**Genotyping population in EGCUT**

Of all the studied biobank participants 33,155 have been genotyped using the Global Screening Array, 8137 HumanOmniExpress beadchip, 2640 HumanCNV370-Duo BeadChips and 6861 Infinium CoreExome-24 BeadChips from Illumina. Furthermore, of 2056 individuals' whole genomes have been sequenced at the Genomics Platform of the Broad Institute.

Sequenced reads were aligned against the GRCh37/hg19 version of the human genome reference using BWA-MEM1 v0.7.7; PCR duplicates were marked using Picard (http://broadinstitute.github.io/picard) v1.136, and the Genome Analysis Toolkit (GATK) v3.4-46 applied for further processing of BAM files and genotype calling. All insertion-deletions (indels) in the Variant Call Format (VCF) were normalized and multiallelic sites split using bcftools (https://samtools.github.io/bcftools/bcftools.html). The following genotypes were set to missing: genotype quality <20, read depth >200, allele balance <0.2 or >0.8 for heterozygous calls. The GATK's Variant Quality Score Recalibration (VQSR) metric was used to filter variants with a truth sensitivity of 99.8% for SNVs and of 99.9% for indels. Furthermore, variants with inbreeding coefficient <-0.3, quality by depth <2 for SNVs and <3 for indels, call rate < 95%, or Hardy-Weinberg equilibrium (HWE) P-value $<1\times10^{-6}$ were excluded.

The genotype calling for the Illumina microarrays was performed using Illumina's GenomeStudio V2010.3 software. The genotype calls for rare variants on the GSA array were corrected using the zCall software (version May 8th, 2012). After variant calling, the data was filtered using PLINK (v.1.90) by sample (call rate >95%, no sex mismatches between phenotype and genotype data, heterozygosity < mean +-3 SE) and marker-wise (HWE p-value $>1\times10^{-6}$, call rate >95%, and for the GSA array additionally by Illumina GenomeStudio GenTrain score >0.6, Cluster Separation Score >0.4). Before the imputation, variants with MAF <1% and C/G or T/A polymorphisms as well as indels were removed, as these genotype calls do not allow precise phasing and imputation. The genotype data obtained on all of the arrays were separately phased using Eagle2 (v. 2.3) and imputed using the BEAGLE (v. 4.1) software implementing a joint Estonian and Finnish reference panel.[17]

**Study outcomes and definitions in EGCUT**

We tested the associations of rs7209826 and rs188810925 with fracture (ICD-10 codes S52.5, S82.6, S22.3, S42.2, S52.6, S22.4, S42.0, S82.8, S72.0, S71.1, S52.1, S32.0, S52.0, S82.4, S82.3, S72.1, S82.5, S22.0, S82.1, S82.7, S82.2, S82.0, S32.5, S32.2, S42.3, S52.2, S52.3, S42.4, S72.3, S52.8, S22.2, S52.4, S42.1, S72.2, S72.4, S32.1, S22.1, S12.2, S32.7, S32.8, S32.4, S82.9, S32.3, S52.9, S12.1, S42.8, S12.7, S72.8, S42.7, S72.9, S22.5, S72.7, S12.0, S42.9), osteoporosis (ICD-10 codes M80 and M81), the prevalent coronary artery disease (ICD-10 codes I20, I21, I22, I23, I24, I25), infarction (ICD-10 codes I21, I22, I25.2) and systolic blood pressure (measured at participant recruitment). For all of the outcome variables with an exception of cardiovascular disease we considered prevalent case statuses reported at recruitment and individuals with records of diagnosis codes reported in the electronic registries before the recruitment. For the outcome of cardiovascular disease, we considered only prevalent cases reported at recruitment.

**Association analyses in EGCUT**

We calculated the effect estimates for the two SNPs using logistic regression and adjusting for age, sex and 15 principal components. All associations were tested under an additive model using glm function with R software (3.3.2).

**Ethical considerations in EGCUT**

Analyses in EGCUT were approved by the Ethics Review Committee of the University of Tartu (243T-12).

**China Kadoorie Biobank (CKB)**
**Study population in CKB**
The China Kadoorie Biobank is a prospective cohort of 512,713 adults aged 30-79 years. Individuals were recruited between the years of 2004-2008 from 5 urban and 5 rural regions across China, as previously described.[18] Baseline information was collected via detailed questionnaire (including demographic/ lifestyle factors and medical history) and physical measurements (which included anthropometry, blood pressure and spirometry). A non-fasting blood sample was taken and separated into plasma and buffy-coat fractions for long-term storage. Long-term follow up is through electronic linkage of each participant's unique national identification number to the Chinese national health insurance system, and established regional registries for death and disease. Health insurance reports include detailed information (e.g. disease description, International Statistical Classification of Diseases and Related Health Problems, 10th Revision [ICD-10] code, and procedure or examination codes) about each hospital admission. Vascular disease events have been reviewed and standardised by clinicians.

**Genotyping population in CKB**
102,783 CKB participants were genotyped using 2 custom-designed Affymetrix Axiom arrays including up to 800K variants, optimised for genome-wide coverage in Chinese populations. Stringent quality control included SNP call rate>0.98, plate effect $p>10^{-6}$, batch effect $p>10^{-6}$, HWE $p>10^{-6}$ (combined 10df Chi-Sq test from 10 regions), biallelic, MAF difference from 1KGP EAS < 0.2, sample call rate >0.95, heterozygosity<mean+3SD, no chrXY aneuploidy, genetically-determined sex concordant with database, and exclusion of recent immigrants to each study area as identified by region-specific principal component analysis, resulting in genotypes for 532,415 variants present on both array versions in 94,592 individuals. Imputation into the 1,000 Genomes Phase 3 reference (EAS MAF>0) using SHAPEIT version 3 and IMPUTE version 4 yielded genotypes for 10,276,634 variants with MAF >0.005 and info >0.3. rs188810925 was not imputed (1KGP EAS MAF=0), rs7209826 was imputed with info=0.986, MAF=0.30.

**Study outcomes and definitions in CKB**
Outcomes, phenotypes and transformations are defined in Table S5. Disease endpoints are for hospitalisations, from electronic linkage to the national health insurance system. Phenotypes were measured either at baseline or in a subset of individuals at the second resurvey of ~5% of the CKB cohort.

**Association analyses in CKB**
To avoid ascertainment biases, unless otherwise specified, controls for binary endpoints were restricted to a 72,795 subset who had been randomly-selected for genotyping (~28k of genotyped samples were selected on the basis of incident CVD or COPD disease events).

Covariates were as specified in Table S5. All analyses used release version 15 of the CKB database and were performed using SAS software (version 9.3; SAS Institute, Inc).

**Ethical considerations in CKB**
Ethical approval for CKB was obtained jointly from the University of Oxford, the Chinese Centre for Disease Control and Prevention (CCDC) and the regional CCDC from the 10 study areas.

**Genome-wide association study (GWAS) consortia**

We supplemented data from UKBB with summary-level data from 9 genome-wide association study (GWAS) consortia, including data for BMD (both ultrasound-derived estimated heel-bone BMD[6] (eBMD) and dual-energy x-ray absorptiometry (DXA)-derived BMD measured at various anatomical sites[8]), coronary heart disease (CHD) and myocardial infarction,[19] stroke,[20] atrial fibrillation,[21] type 2 diabetes mellitus[22] and glycaemic traits,[23,24] serum lipid fractions,[25] anthropometric traits,[12,13] and chronic kidney disease.[26,27] Where available, we selected data pertaining to analyses conducted in European-ancestry individuals. Further details for each consortium are provided in Table S2.

**Selection of proxy SNPs**

Since the selected instrument SNPs (rs7209826 and rs188810925) were not available in all GWAS consortia datasets, we identified SNPs to be used as proxies, using HaploReg.[28] We set an $r^2$ threshold of >0.9 (in European ancestry individuals) for selection of suitable proxies. We identified 14 SNPs to be in high LD ($r^2$=0.99) with rs7209826. Of these 14, we selected rs7220711 as a proxy for rs7209826 based on high LD ($r^2$=0.99 in European ancestry populations), availability across most consortia, and prior functional evidence linking rs7220711 to *SOST* expression.[29] In addition, we validated the effect of rs7220711 on various measures of BMD as being comparable to that of rs7209826 (Figure S2). There were no suitable proxies ($r^2$>0.9) for rs188810925.

**Scaling of allelic estimates**

We scaled allelic estimates pertaining to risk of osteoporosis, fractures, cardiometabolic outcomes and quantitative traits to an increase in BMD equivalent to that reported in a phase II RCT of 12 months of 210mg romosozumab monthly.[30] This corresponds to the dose evaluated in phase III RCTs of romosozumab, and represents a 0.09 g/cm$^2$ increase in lumbar spine BMD (LS-BMD) in postmenopausal women.[30,31] We selected lumbar spine BMD as the reference phenotype as this was the only BMD phenotype for which we had genetic data[8] and clinical trial data[31] in the same units of measurements (i.e. g/cm2). To do this, we derived the allelic effect estimates (in g/cm$^2$ units) for the association of rs7209826 and rs188810925 with LS-BMD from a previous large-scale GWAS of LS-BMD[8] (0.008 and 0.016 g/cm$^2$, respectively). We then derived a scaling factor for each SNP by dividing the increase in LS-BMD arising from treatment with 210mg romosozumab monthly for 12 months (0.09 g/cm$^2$) by the allelic effect estimate for LS-BMD. All per-allele effect estimates (log(OR) and the standard error of log(OR) for binary outcomes and beta and the standard error of beta for quantitative traits) were subsequently multiplied by these scaling factors to estimate the effects expected with 12 months of 210mg romosozumab monthly

**Meta-analysis of scaled allelic estimates**

The R-package METAFOR was used for performing all meta-analyses of scaled allelic estimates.[32] Fixed-effect meta-analyses were used in all instances. Cochran's Q test was used to evaluate heterogeneity for all meta-analyses performed (see Table S10).

**Identification of phase III clinical trials of sclerostin inhibitors**

We searched for all phase III RCTs performed for sclerostin inhibitors. We collected a record of trials conducted for sclerostin inhibitors from the websites of the agents' developers, and supplemented this with further searches on clinical trials registries (ClinicalTrials.gov, EU Clinical Trials Register and International Clinical Trials Registry Platform) and PubMed, using the key-words ("sclerostin" OR "romosozumab" OR "AMG-785") AND ("trial" OR "randomized controlled trial" OR "RCT" OR "randomised controlled trial").

**Meta-analysis of randomized controlled trials**

For meta-analysis of cardiovascular events (using data from the FRAME[33], ARCH[33] and BRIDGE[34] trials; Table S6), we used the Mantel-Haenszel method without continuity correction. This method has been shown to perform relatively well when events are rare, and is also the default fixed-effect meta-analysis method recommended by the Cochrane collaboration.[35–37] We performed additional sensitivity analyses using the Peto method, a fixed-effect method shown to provide relatively unbiased results if within-trial intervention and control groups are of approximately equal size and if the effect size is modest.[35,36] Meta-analysis of fracture data was performed using inverse variance weighted fixed-effect meta-analysis of fracture risk at 12 months in the FRAME[38] and ARCH[33] trials (Table S12). All meta-analyses of trial data were performed using the R-package METAFOR. Cochran's Q test was used to evaluate heterogeneity for all meta-analyses of trial data (Table S11).

#### Supplementary Figures

##### Figure S1. Overview of study design.

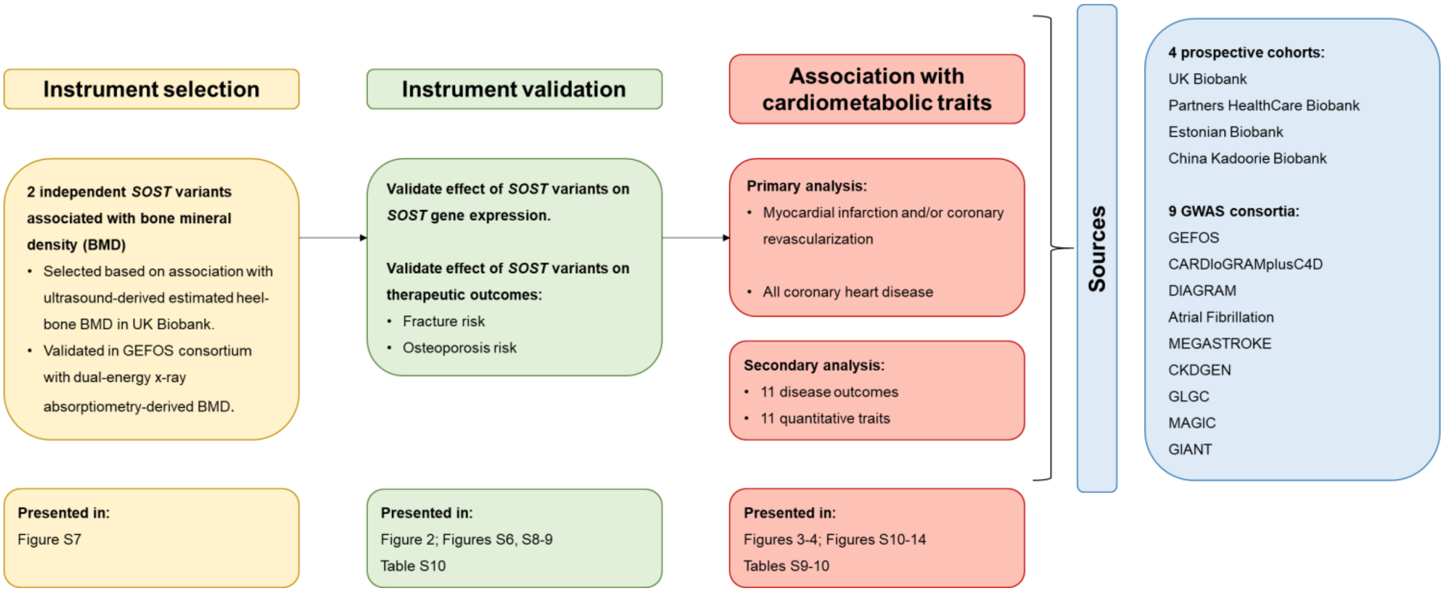


We selected genetic variants in the *SOST* locus associated with bone mineral density. We validated the effect of these variants on therapeutic outcomes of interest for therapeutic sclerostin inhibition, i.e. risk of fracture and osteoporosis. We followed this with two sets of analyses. In the primary analysis, we examined the effect of the *SOST* variants on myocardial infarction and/or coronary revascularization and on all coronary heart disease. In the secondary analysis, we studied the association of the *SOST* variants with a further 11 cardiometabolic outcomes and 11 quantitative traits. Data were sourced from 4 prospective cohorts and 9 GWAS consortia. BMD, bone mineral density; CARDIoGRAMplusC4D, Coronary Artery Disease Genome wide Replication and Meta-analysis (CARDIoGRAM) plus The Coronary Artery Disease (C4D) Genetics; CKDGEN, Chronic Kidney Disease Genetics; DIAGRAM, Diabetes Genetics Replication and Meta-analysis; GEFOS, Genetic Factors for Osteoporosis Consortium; GIANT, Genetic Investigation of Anthropometric Traits; GLGC, Global Lipid Genetics Consortium; MAGIC, Meta-Analysis of Glucose and Insulin-related traits Consortium.

**Figure S2. Regional association plots of *SOST* locus in GWAS of heel-bone estimated bone mineral density in UKBB (n = 142,487).**

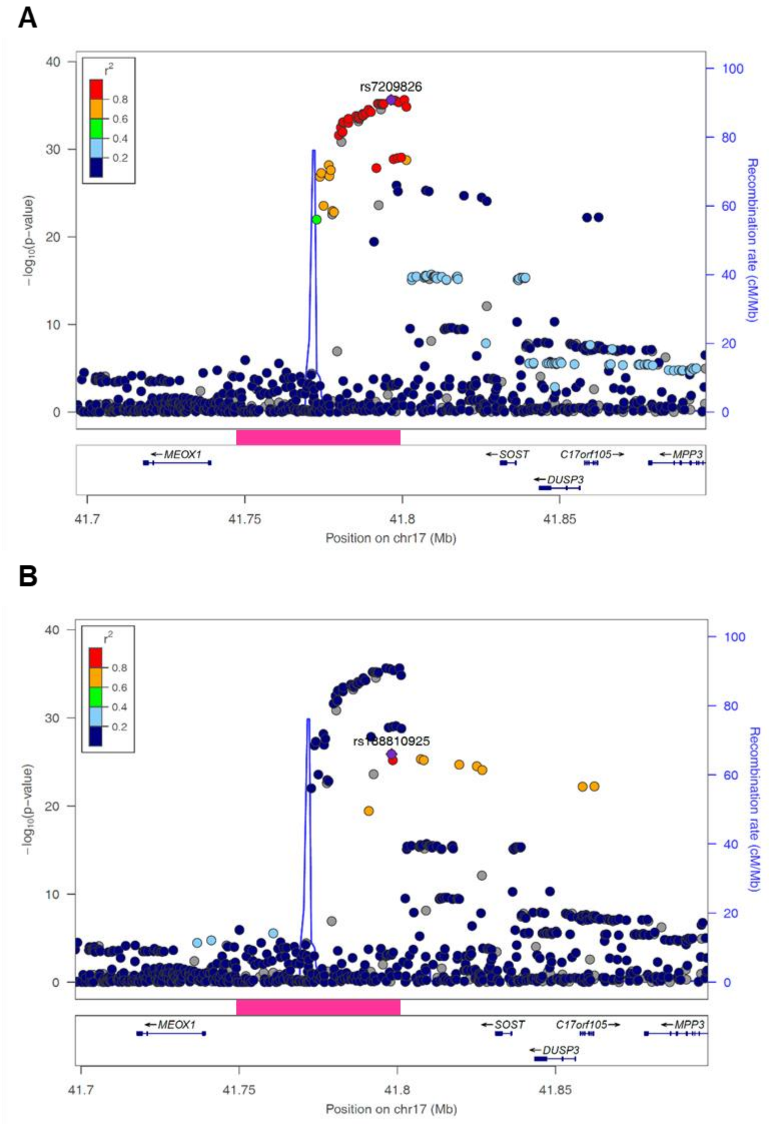


Two independent ($r^2$=0.13 in UKBB) genome-wide significant variants were identified in the *SOST* locus, rs188810925 and rs7209826. Each plotted point represents a SNP; colours of the plotted points indicate the linkage disequilibrium (in $r^2$) of each SNP with the instrument SNPs (indicated as purple points; rs7209826 (Panel A) and rs188810925 (Panel B)). The y-axis indicates the $-\log_{10}$(p-value for association with bone mineral density) for each plotted point, with chromosomal coordinates (and gene locations) shown on the x-axis. The magenta block indicates the approximate location of the 52kb deletion area associated with van Buchem disease, a Mendelian disorder of decreased *SOST* expression leading to low levels of sclerostin and a bone overgrowth phenotype. GWAS data sourced from GEFOS consortium.[5]

**Figure S3. Per-allele associations of rs7209826 and selected proxy (rs7220711; $r^2 = 0.99$) with various bone mineral density (BMD) measures.**

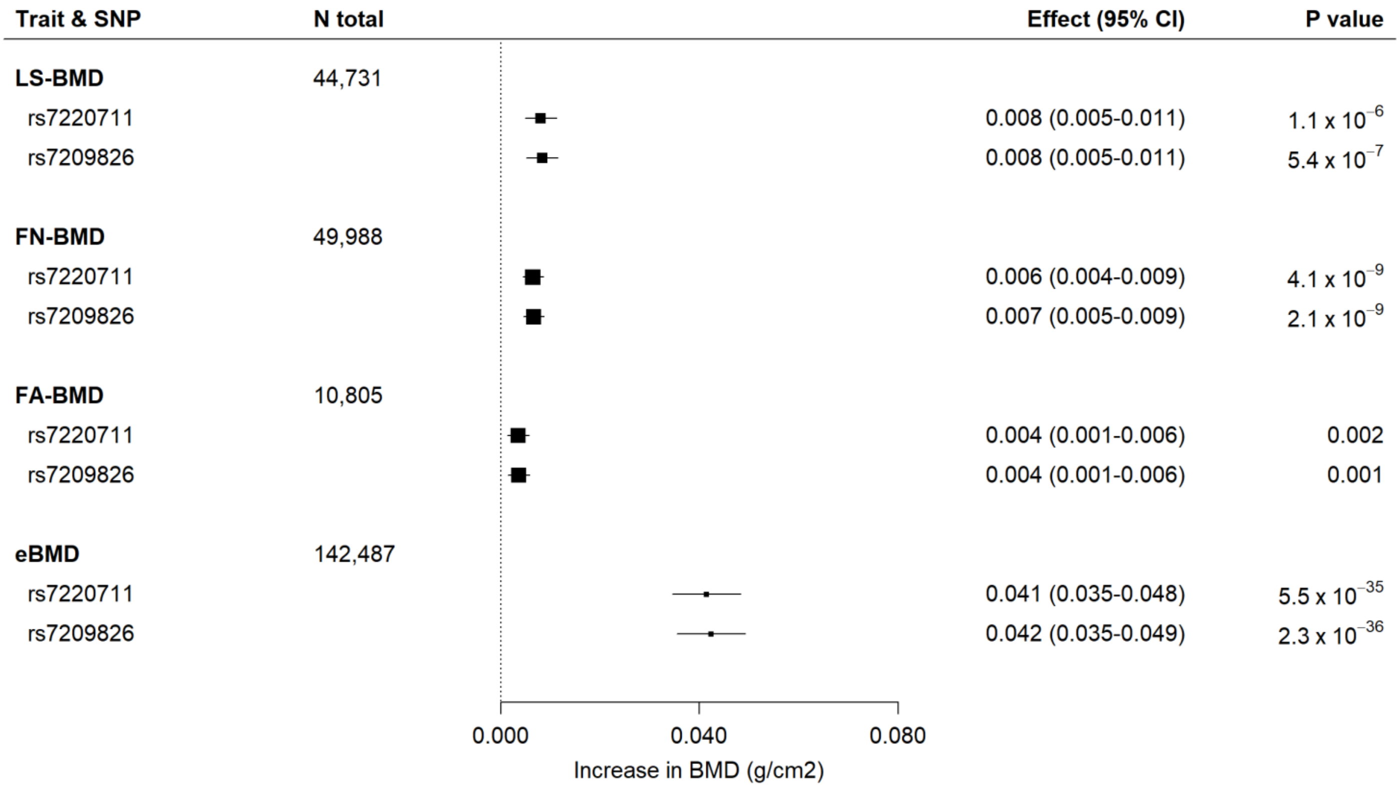


All data are plotted in $g/cm^2$ units and effects are per minor allele. Measures for FN-BMD, LS-BMD, FA-BMD were derived by dual energy X-ray absorptiometry (DXA); measures for eBMD were derived by heel-bone ultrasound. Boxes represent point estimates of effects. Lines represent 95% confidence intervals. LS-BMD, Lumbar Spine BMD; FN-BMD, Femoral Neck BMD; FA-BMD, Forearm BMD; eBMD, estimated heel-bone BMD. Data sourced from GEFOS consortium.[5,6]

**Figure S4. Meta-analysis of romosozumab and risk of cardiovascular events from phase III randomized controlled trials, using Peto Method.**

| Endpoint & Trial | Romosozumab Events | Romosozumab Total | Comparator Events | Comparator Total | OR (95% CI) | P value |
|---|---|---|---|---|---|---|
| **Cardiac ischemic events** | | | | | | |
| ARCH | 16 | 2040 | 6 | 2014 | 2.46 (1.07-5.69) | |
| BRIDGE | 3 | 163 | 0 | 81 | 2.58 (0.32-20.92) | |
| **Overall** | **19** | **2203** | **6** | **2095** | **2.63 (1.19-5.80)** | **0.02** |
| **Cerebrovascular events** | | | | | | |
| ARCH | 16 | 2040 | 7 | 2014 | 2.17 (0.96-4.92) | |
| BRIDGE | 3 | 163 | 1 | 81 | 1.45 (0.18-11.80) | |
| **Overall** | **19** | **2203** | **8** | **2095** | **2.06 (0.96-4.41)** | **0.06** |
| **Serious cardiovascular events** | | | | | | |
| FRAME | 44 | 3581 | 41 | 3576 | 1.07 (0.70-1.64) | |
| ARCH | 50 | 2040 | 38 | 2014 | 1.30 (0.85-1.99) | |
| BRIDGE | 8 | 163 | 2 | 81 | 1.86 (0.49-7.09) | |
| **Overall** | **102** | **5784** | **81** | **5671** | **1.21 (0.90-1.62)** | **0.20** |

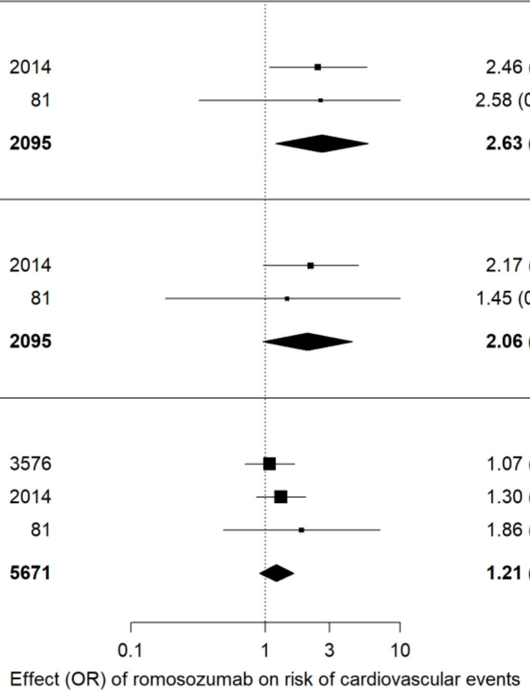


*Events* represents number of events adjudicated per arm, during initial 12-month double-blind period in each trial. The *romosozumab*-group received 210mg romosozumab monthly in all trials; *comparator*-group received placebo (FRAME and BRIDGE trials) or alendronate (ARCH). Estimates derived using the Peto method. Outcome data for *cardiac ischemic events* and *cerebrovascular events* were only available for the ARCH and BRIDGE trials. Boxes represent point estimates of effects. Lines represent 95% confidence intervals. OR, odds ratio; CI, confidence interval.

**Figure S5.** ***SOST*** **expression is highest in arterial tissues.**

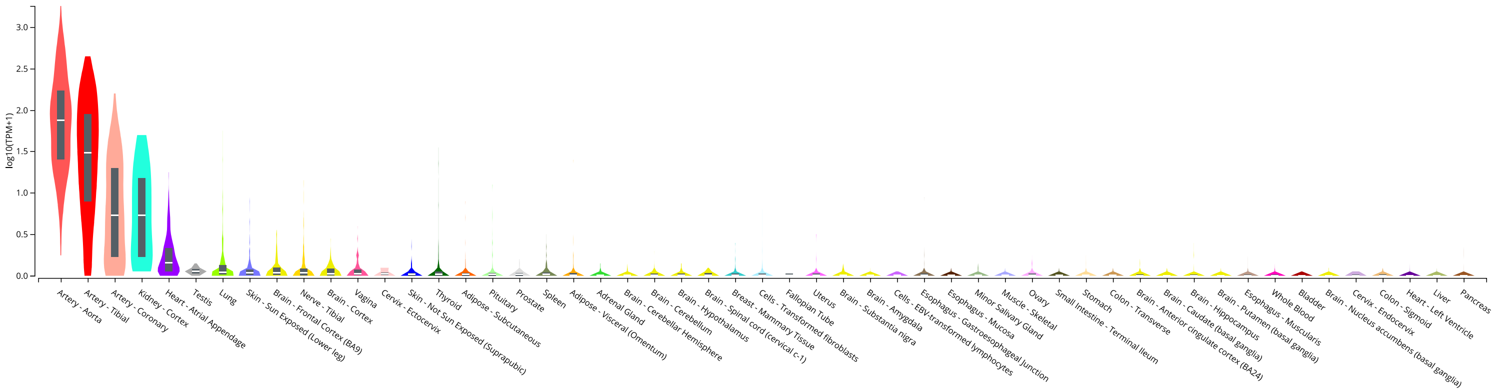


Expression values are shown as $\log_{10}$(transcripts per million). Tissues are ranked from highest to lowest expression level across 53 tissue types (bone tissue not available in dataset). Box plots represent median and $25^{th}$/$75^{th}$ percentiles. Sourced from GTEx Analysis Release V7. TPM = transcripts per million.

**Figure S6.** ***SOST*** **mRNA expression by rs188810925 and rs7209826 genotype.**

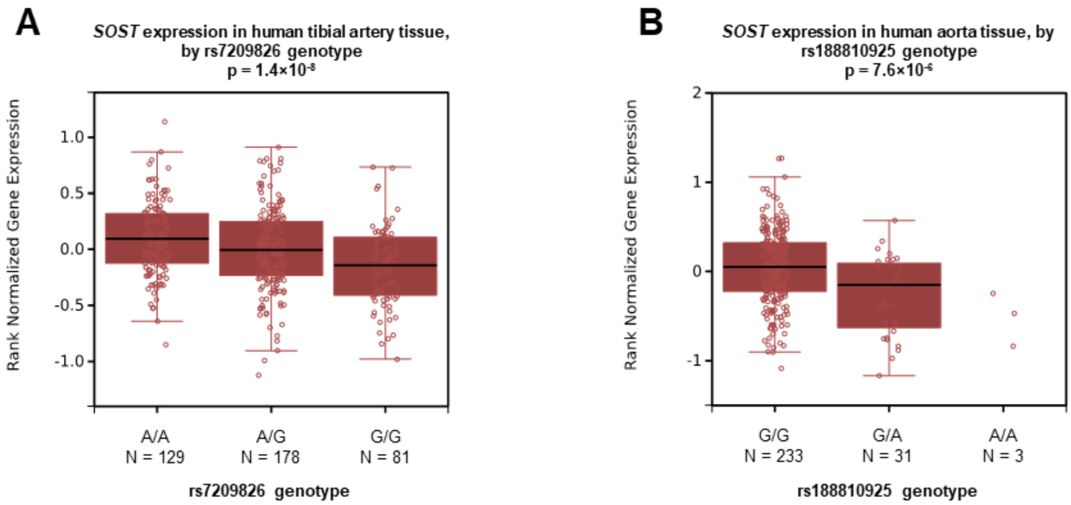


Shown are the strongest associations, across 53 tissue types, for rs7209826 (human tibial artery; Panel A) and rs188810925 (human aorta; Panel B). The minor alleles for both variants associate with lower *SOST* expression. Sourced from GTEx Analysis Release V7.

**Figure S7. Per-allele associations of rs7209826 and rs188810925 with various bone mineral density (BMD) measures.**

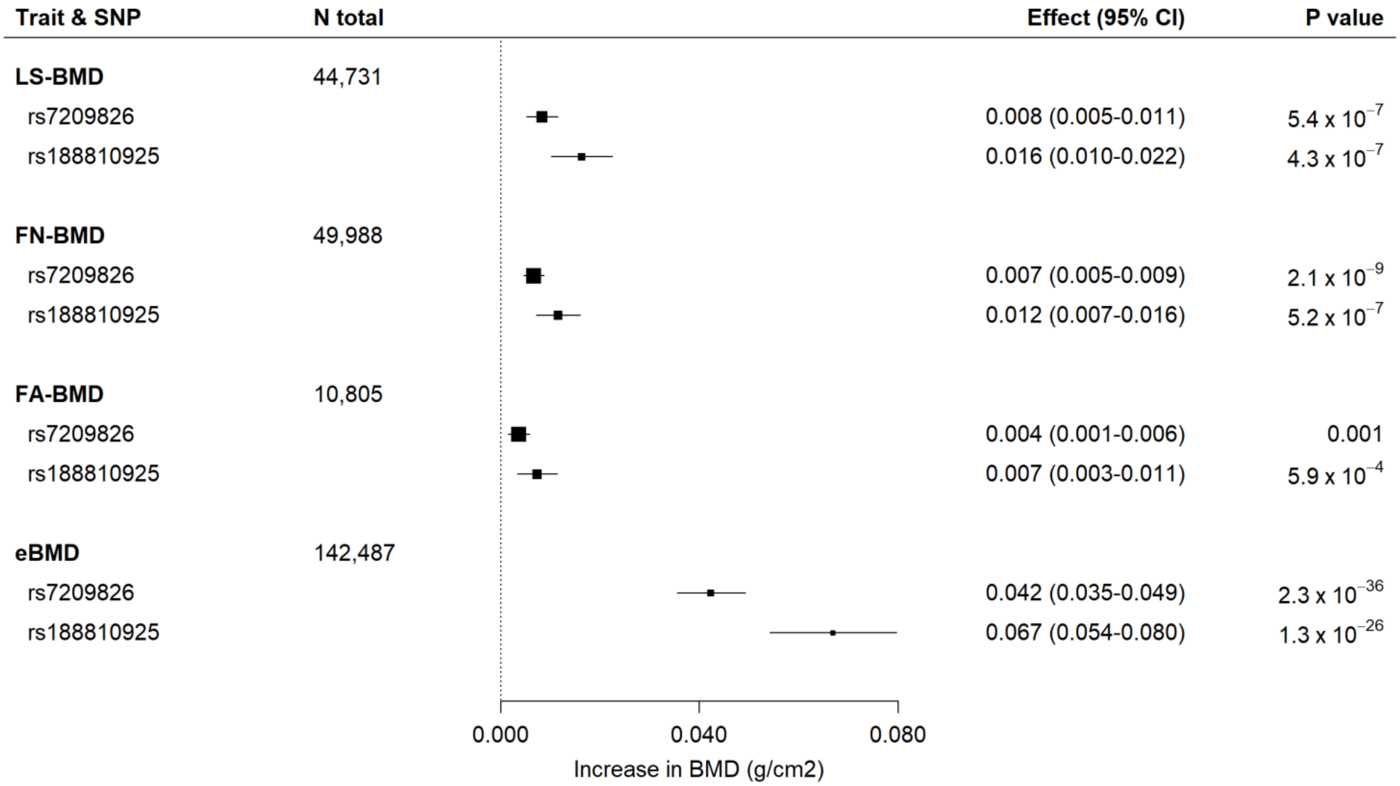


All data are plotted in g/cm$^2$ units and effects are per minor allele. Measures for FN-BMD, LS-BMD, FA-BMD were derived by dual energy X-ray absorptiometry (DXA); measures for eBMD were derived by heel-bone ultrasound. Boxes represent point estimates of effects. Lines represent 95% confidence intervals. LS-BMD, Lumbar Spine BMD; FN-BMD, Femoral Neck BMD; FA-BMD, Forearm BMD; eBMD, estimated heel-bone BMD. Data sourced from GEFOS consortium.[5,6]

**Figure S8. Study-specific scaled estimates and meta-analysis of BMD-increasing *SOST* variants with risk of osteoporosis and fracture.**

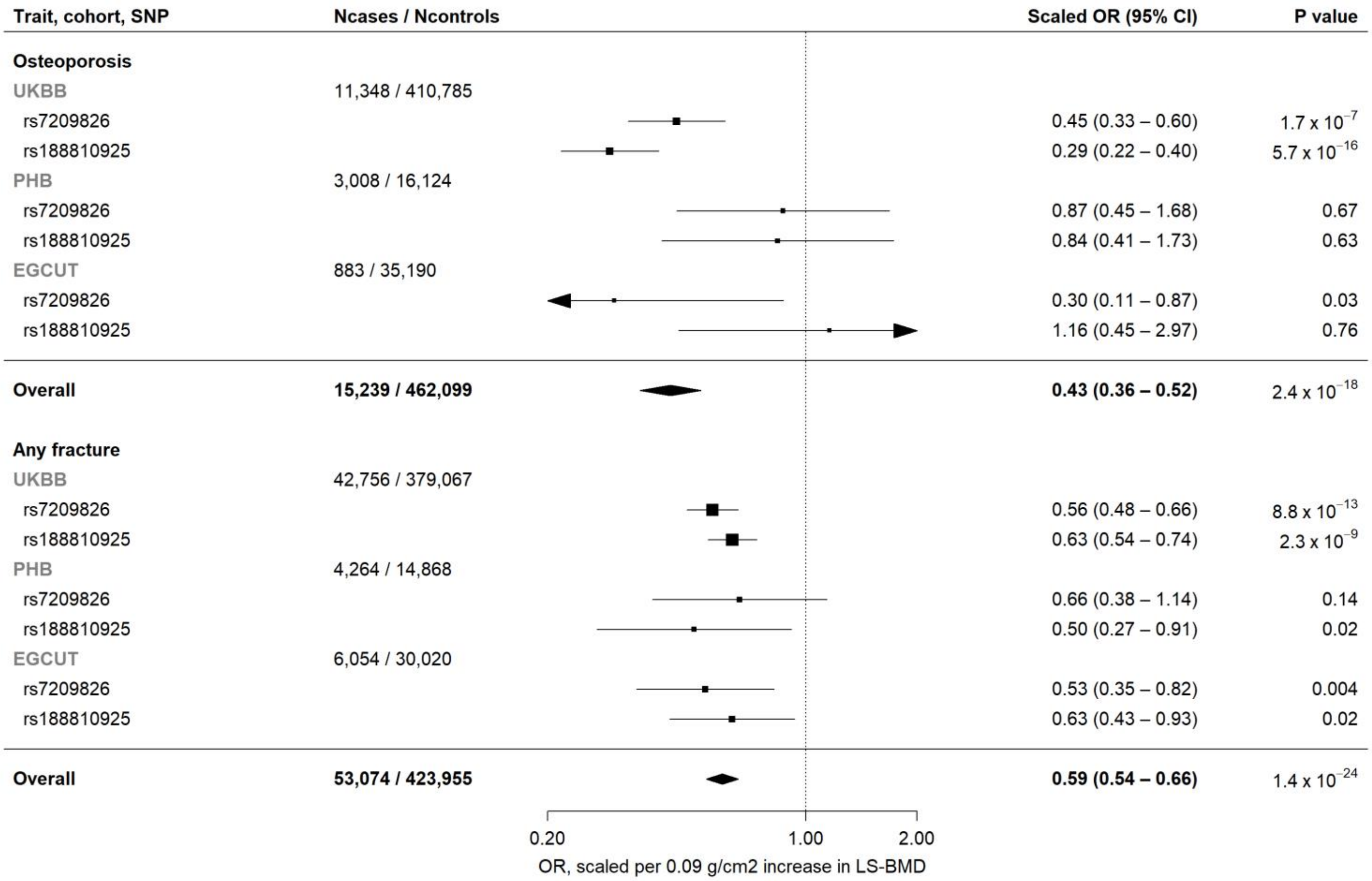


Estimates are scaled to match the effect of 210mg romosozumab monthly for 12 months on lumbar spine bone mineral density (0.09 g/cm$^2$; see Methods) and aligned to the BMD-increasing alleles. Boxes represent point estimates of effects. Lines represent 95% confidence intervals. UKBB, UK Biobank; PHB, Partners HealthCare Biobank; EGCUT, Estonian Biobank; OR, odds ratio; CI, confidence interval.

**Figure S9. Scaled estimates and meta-analysis of BMD-increasing *SOST* variants with risk of fracture in the preceding 5 years, at different anatomical sites, in UKBB.**

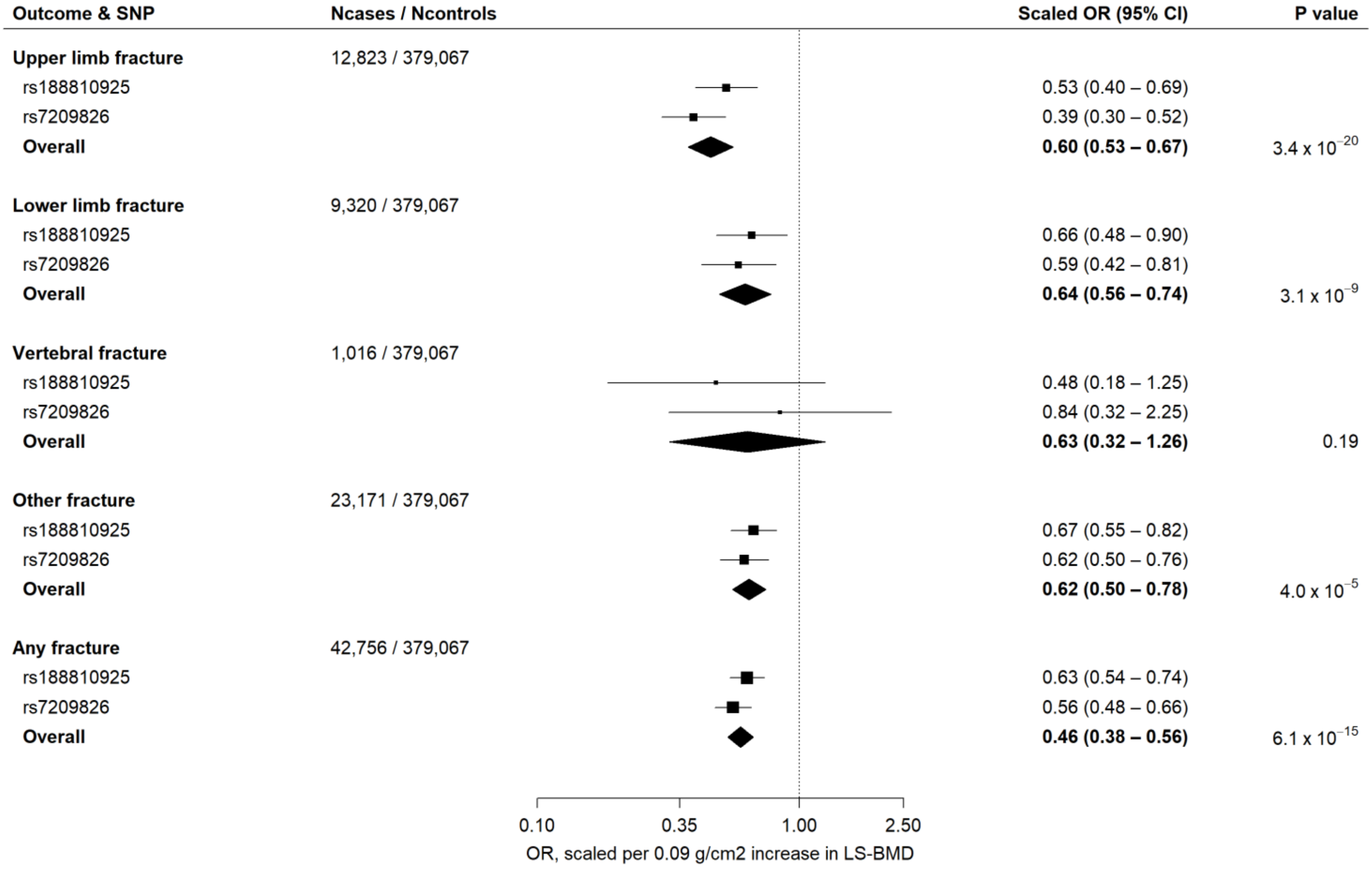


Estimates are scaled to match the effect of 210mg romosozumab monthly for 12 months on lumbar spine bone mineral density (0.09 g/cm$^2$; see Methods) and aligned to the BMD-increasing alleles. *Upper limb fractures* included fractures reported as involving the wrist or arm; *Lower limb fractures* included fractures reported as involving the ankle, leg or hip; *Vertebral fractures* included fractures reported as involving the spine*; Other fractures* included all fractures reported as involving bones other than those specified above. *Any fracture* included any fracture or broken bone sustained in the preceding 5 years. Boxes represent point estimates of effects. Lines represent 95% confidence intervals. OR, odds ratio; CI, confidence interval.

**Figure S10. Study-specific scaled estimates and meta-analysis of BMD-increasing *SOST* variants with risk of myocardial infarction and/or coronary revascularization and coronary heart disease.**

| Trait, cohort, SNP | Ncases / Ncontrols | Scaled OR (95% CI) | P value |
|---|---|---|---|
| **Myocardial Infarction and/or coronary revascularization** | | | |
| UKBB | 21,538 / 387,642 | | |
| rs7209826 | | 1.28 (1.02 – 1.60) | 0.03 |
| rs188810925 | | 1.18 (0.96 – 1.45) | 0.13 |
| PHB | 3,288 / 12,439 | | |
| rs7209826 | | 1.93 (0.97 – 3.85) | 0.06 |
| rs188810925 | | 1.02 (0.48 – 2.14) | 0.97 |
| EGCUT | 1,147 / 28,062 | | |
| rs7209826 | | 0.71 (0.27 – 1.91) | 0.5 |
| rs188810925 | | 0.40 (0.16 – 1.00) | 0.05 |
| CARDIoGRAMplusC4D | 43,676 / 128,199 | | |
| rs7209826 | | 1.04 (0.83 – 1.30) | 0.73 |
| rs188810925 | | 1.36 (1.05 – 1.77) | 0.02 |
| **Overall** | **69,649 / 556,342** | **1.18 (1.06 – 1.32)** | 0.003 |
| **Coronary heart disease** | | | |
| UKBB | 36,078 / 387,642 | | |
| rs7209826 | | 1.27 (1.07 – 1.52) | 0.007 |
| rs188810925 | | 1.14 (0.96 – 1.34) | 0.13 |
| PHB | 6,599 / 12,439 | | |
| rs7209826 | | 1.29 (0.77 – 2.18) | 0.33 |
| rs188810925 | | 0.91 (0.51 – 1.60) | 0.74 |
| EGCUT | 2,851 / 28,062 | | |
| rs7209826 | | 0.75 (0.38 – 1.45) | 0.39 |
| rs188810925 | | 0.45 (0.24 – 0.84) | 0.01 |
| CARDIoGRAMplusC4D | 60,801 / 123,504 | | |
| rs7209826 | | 1.00 (0.82 – 1.22) | 1 |
| rs188810925 | | 1.07 (0.84 – 1.36) | 0.57 |
| **Overall** | **106,329 / 551,647** | **1.10 (1.00 – 1.20)** | 0.04 |

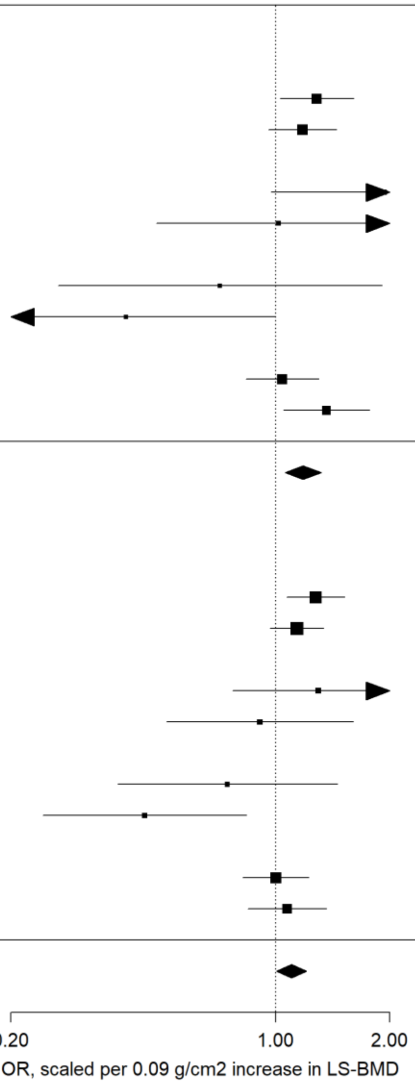


Estimates are scaled to match the effect of 210mg romosozumab monthly for 12 months on lumbar spine bone mineral density (0.09 g/cm$^2$; see Methods) and aligned to the BMD-increasing alleles. Boxes represent point estimates of effects. Lines represent 95% confidence intervals. UKBB, UK Biobank; PHB, Partners HealthCare Biobank; EGCUT, Estonian Biobank; OR, odds ratio; CI, confidence interval.

**Figure S11. Study-specific scaled estimates and meta-analysis of BMD-increasing *SOST* variants with risk of 11 cardiometabolic outcomes.**

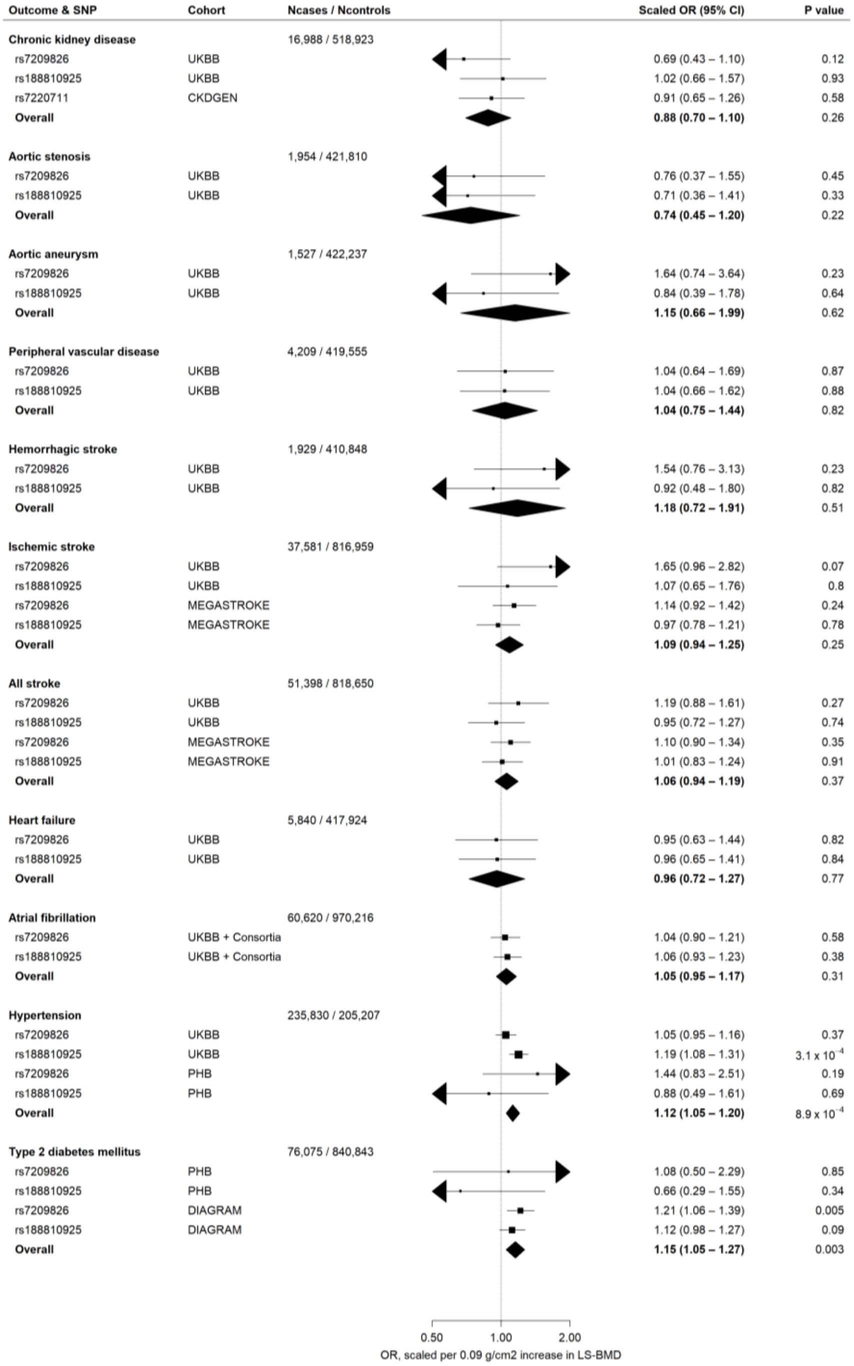


Estimates are scaled to match the effect of 210mg romosozumab monthly for 12 months on lumbar spine bone mineral density (0.09 g/cm$^2$; see Methods) and aligned to the BMD-increasing alleles. Consortia+UKBB implies that the meta-analysed estimate includes data from UK Biobank and from a relevant consortium for the same outcome; AF+ and DIAGRAM+ refers to consortia data which includes all cases from UKBB with cases from various other cohorts. For each meta-analysis, the p-value for heterogeneity was > 0.05 (not shown in plot). The significance threshold was set at 0.0045 (Bonferroni-adjusted; 0.05 / 11). As data from the CKDGEN consortium did not include either rs7209826 or rs188810925, we selected rs7220711 as a proxy for rs7209826 (no suitable proxy was available for rs188810925; see Methods). Boxes represent point estimates of effects. Lines represent 95% confidence intervals. UKBB, UK Biobank; OR, odds ratio; CI, confidence interval; PHB, Partners HealthCare Biobank; EGCUT, Estonian Biobank.

**Figure S12. Study-specific scaled estimates and meta-analysis of BMD-increasing *SOST* variants with systolic and diastolic blood pressure.**

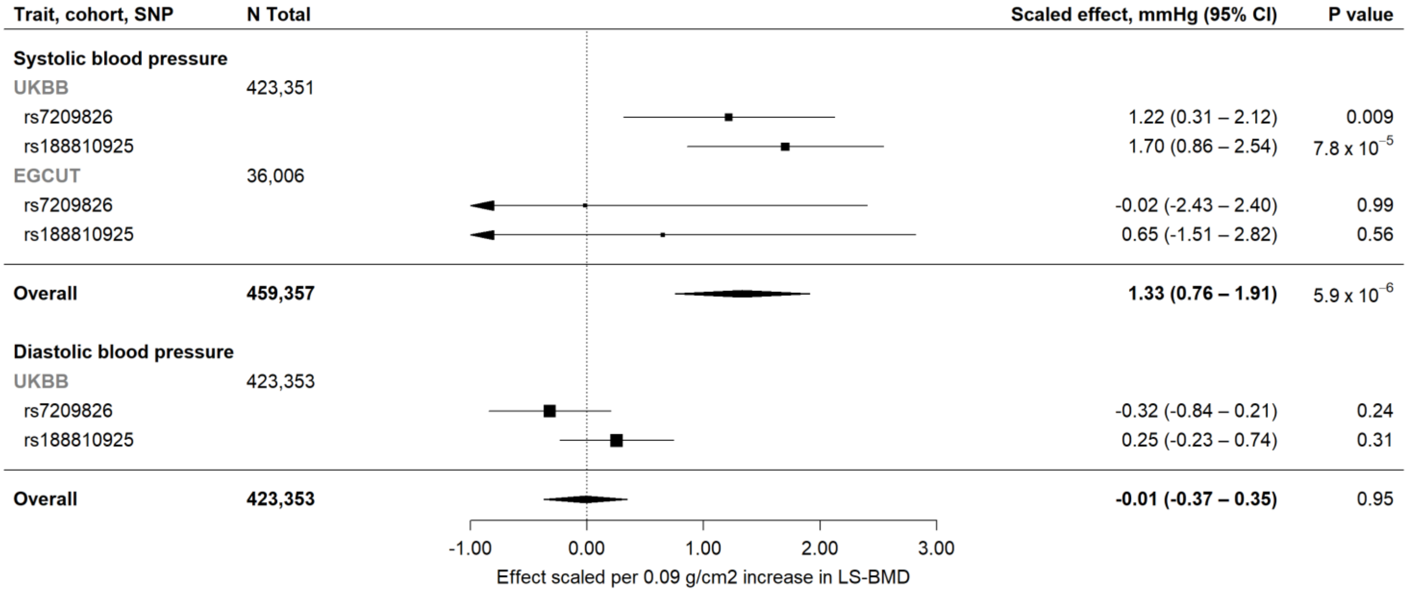


Estimates are scaled to match the effect of 210mg romosozumab monthly for 12 months on lumbar spine bone mineral density (0.09 g/cm$^2$; see Methods) and aligned to the BMD-increasing alleles. Boxes represent point estimates of effects. Lines represent 95% confidence intervals. BP, blood pressure; CI, confidence interval; PHB, Partners HealthCare Biobank; EGCUT, Estonian Biobank.

**Figure S13. Study-specific scaled estimates and meta-analysis of BMD-increasing *SOST* variants with waist to hip ratio (WHR; adjusted for body mass index), body mass index (BMI), LDL cholesterol, HDL cholesterol and triglycerides.**

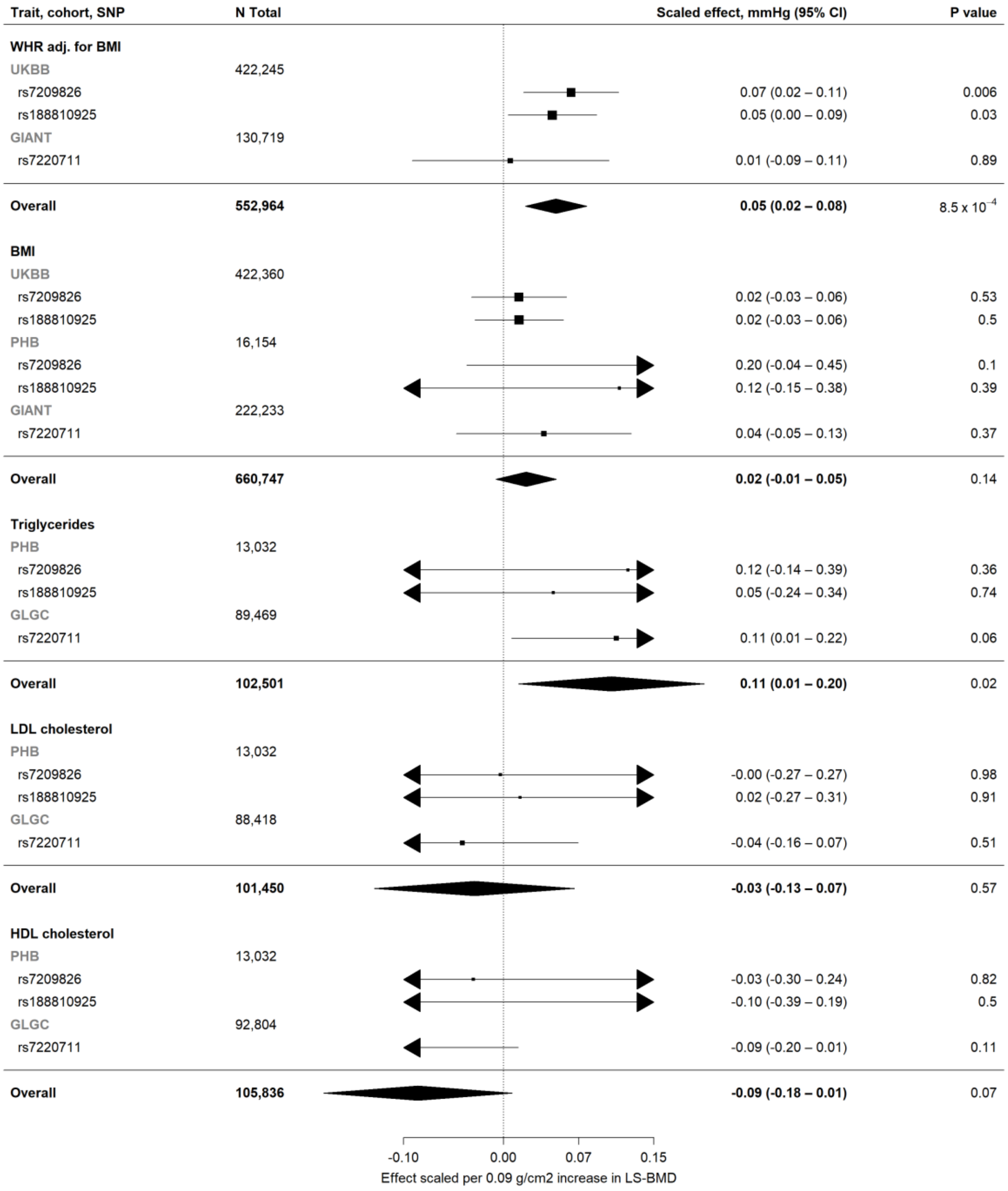


Estimates are scaled to match the effect of 210mg romosozumab monthly for 12 months on lumbar spine bone mineral density (0.09 g/cm$^2$; see Methods) and aligned to the BMD-increasing alleles. As data from the GIANT consortium did not include either rs7209826 or rs188810925, we selected rs7220711 as a proxy for rs7209826 (no suitable proxy was available for rs188810925; see Methods). Boxes represent point estimates of effects in standard deviation units. Lines represent 95% confidence intervals. CI, confidence interval; WHR, waist to hip ratio; adj, adjusted; BMI, body mass index; UKBB, UK Biobank; PHB, Partners HealthCare Biobank; EGCUT, Estonian Biobank.

**Figure S14. Scaled estimates and meta-analysis of BMD-increasing *SOST* variants with glomerular filtration rate estimated from serum creatinine (eGFR) in the CKDGEN consortium.**

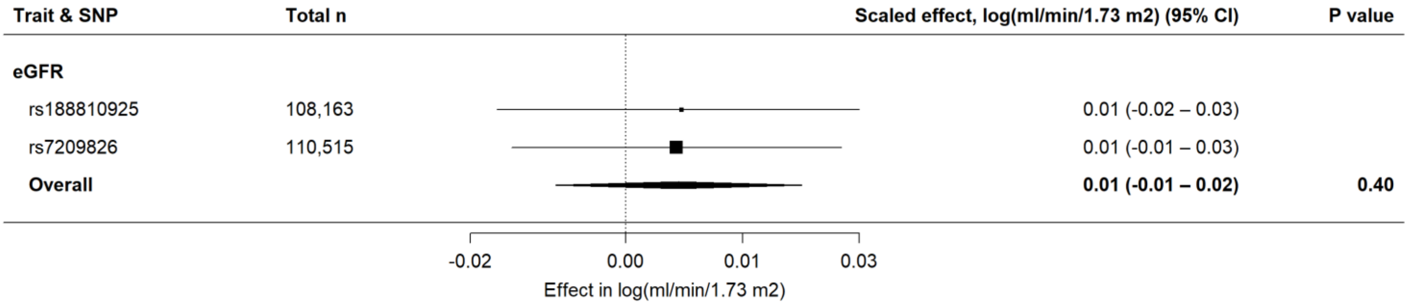


Estimates are scaled to match the effect of 210mg romosozumab monthly for 12 months on lumbar spine bone mineral density (0.09 g/cm$^2$; see Methods) and aligned to the BMD-increasing alleles. Boxes represent point estimates of effects in log(ml/min/1.73 m$^2$) units. Lines represent 95% confidence intervals. eGFR, estimated glomerular filtration rate; CI, confidence interval.

**Figure S15. Per-allele association of rs7209826 with estimated heel-bone bone mineral density (eBMD) in UKBB and China Kadoorie Biobank.**

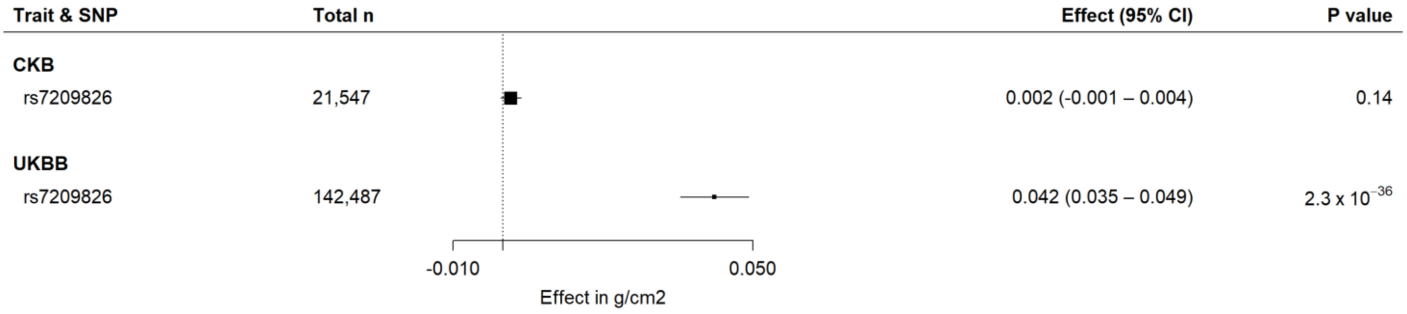


All data are plotted in g/cm$^2$ units and effects are per G-allele. Measures for eBMD were derived by heel-bone ultrasound. Boxes represent point estimates of effects. Lines represent 95% confidence intervals. P-het refers to p-value from Cochran's Q test. eBMD, estimated heel-bone BMD; UKBB, UK Biobank. UKBB data sourced from GEFOS consortium.[5]

**Figure S16. Regional association plot of *SOST* locus in GWAS of heel-bone estimated bone mineral density in China Kadoorie Biobank (n = 21,547).**

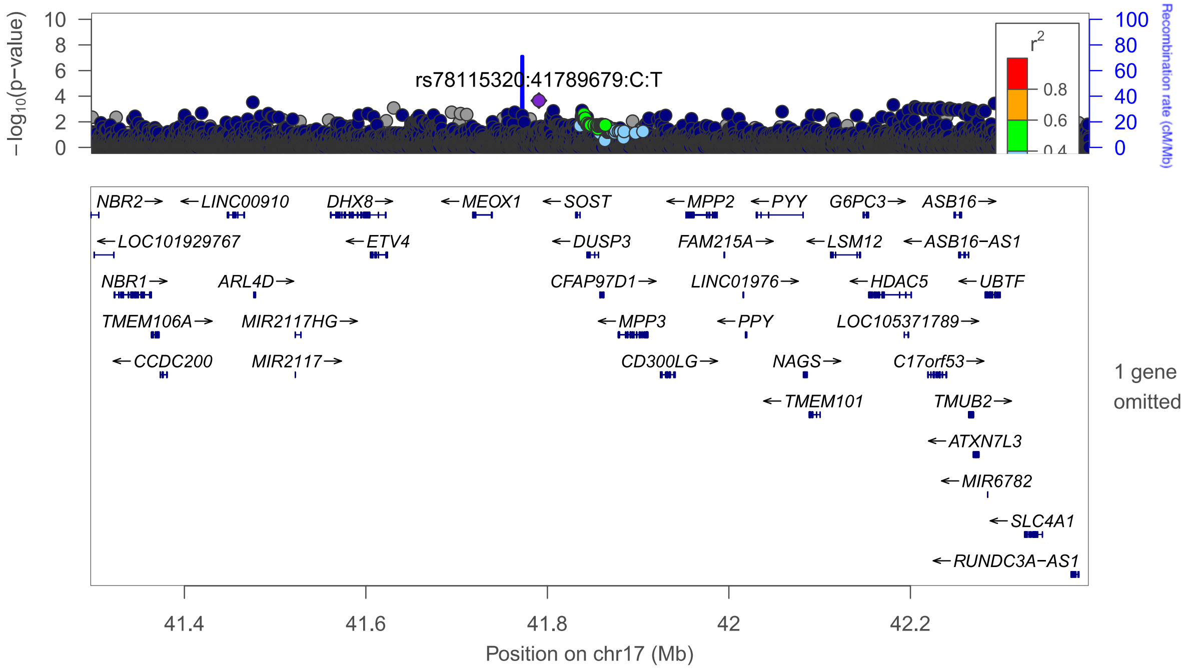


Locus plot showing no evidence of a genome-wide significant genetic association signal in the *SOST* locus in China Kadoorie Biobank. Each plotted point represents a SNP; colours of the plotted points indicate the linkage disequilibrium (in $r^2$) of each SNP with the lead SNP (rs78115320 in this plot). The y-axis indicates the $-\log_{10}$(p-value for association with bone mineral density) for each plotted point, with chromosomal coordinates (and gene locations) shown on the x-axis.

**Figure S17. Meta-analysis of romosozumab and risk of clinical fracture from phase III randomized controlled trials.**

| | Romosozumab | | Comparator | | | |
|---|---|---|---|---|---|---|
| Outcome & Trial | Events | Total | Events | Total | HR (95% CI) | P value |
| **Clinical fracture** | | | | | | |
| FRAME | 58 | 3589 | 90 | 3591 | 0.64 (0.46-0.89) | |
| ARCH | 79 | 2046 | 110 | 2047 | 0.72 (0.54-0.96) | |
| **Overall** | **137** | **5635** | **200** | **5638** | **0.68 (0.55-0.85)** | **0.0006** |

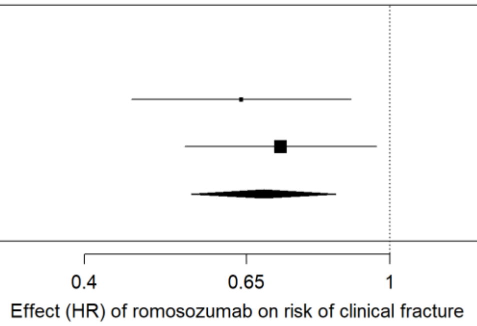


*Clinical fracture* represents a composite of nonvertebral fracture and symptomatic vertebral fracture. *Events* represents number of events adjudicated per arm, during initial 12-month double-blind period in each trial. The *romosozumab*-group received 210mg romosozumab monthly in both trials; *comparator*-group received placebo (FRAME) or alendronate (ARCH). Estimates derived using fixed-effects meta-analysis. Boxes represent point estimates of effects. Lines represent 95% confidence intervals. OR, odds ratio; CI, confidence interval.

**Supplementary Tables**

**Table S1. Baseline characteristics in UK Biobank.**

| **Variable** | |
|---|---|
| Individuals | 423,761 |
| Female, no (%) | 228,628 (54%) |
| Age, mean (SD), years | 56.8 (7.9) |
| BMI, mean (SD), kg/m2 | 27.4 (4.8) |
| Systolic Blood Pressure (SD), mmHg | 141.2 (20.7) |
| Diastolic Blood Pressure (SD), mmHg | 84.3 (11.3) |
| Coronary Heart Disease, N (%) | 36,078 (8.5%) |

SD, standard deviation.

**Table S2. Genome-wide association studies and consortia included in present study.**

| Consortium | Trait(s) or outcome(s) analyzed for present study | PMID | Cases (or total) | Controls |
|---|---|---|---|---|
| GEFOS | eBMD | 28869591 | 142,487 | NA |
| | LS-BMD | 26367794 | 44,731 | NA |
| | FN-BMD | 26367794 | 49,988 | NA |
| | FA-BMD | 26367794 | 10,805 | NA |
| DIAGRAM* | Type 2 Diabetes Mellitus (non-BMI adjusted) | 30297969 | 74,124 | 824,006 |
| Atrial fibrillation* | Atrial fibrillation | 30061737 | 60,620 | 970,216 |
| CARDIoGRAMplusC4D | All CHD | 26343387 | 60,801 | 123,504 |
| | Myocardial infarction | 26343387 | 43,676 | 128,199 |
| MEGASTROKE | Ischemic stroke | 29531354 | 34,217 | 406,111 |
| | Any stroke | 29531354 | 40,585 | 406,111 |
| CKDGEN | eGFR (based on serum creatinine) | 28452372 | 110,515 | NA |
| | Chronic kidney disease | 26831199 | 12,385 | 104,780 |
| GLGC | High-density lipoprotein cholesterol | 24097068 | 92,804 | NA |
| | Low-density lipoprotein cholesterol | 24097068 | 88,418 | NA |
| | Triglycerides | 24097068 | 89,469 | NA |
| MAGIC | HbA1c | 28898252 | 123,665 | NA |
| | Fasting glucose | 22581228 | 58,074 | NA |
| | Fasting insulin | 22581228 | 51,750 | NA |
| GIANT | BMI | 25673413 | 222,233 | NA |
| | Waist-hip ratio, adjusted for BMI | 25673412 | 130,719 | NA |

PMID refers to PubMed identification number of the principal publication within which the data-source was first reported. "NA" is used for quantitative traits. None of these data-sources included UKBB data, with the exception of the data-sources marked with * (DIAGRAM and Atrial Fibrillation), which included all cases identified in UKBB and further cases from other cohorts. CHD, coronary heart disease; eBMD, estimated heel-bone bone mineral density; LS-BMD, lumbar-spine bone mineral density; FN, femoral neck BMD; FA-BMD, forearm BMD; BMI, body mass index; UKBB, UK Biobank; eGFR, estimated glomerular filtration rate; HbA1c, glycated haemoglobin.

### Table S3. Definitions of outcomes analysed in UK Biobank.

| Outcome | ICD-10 | ICD-9 | OPCS-4 | Nurse Interview, Non-cancer | NI, operation | Self-reported Diagnosis by Doctor | Description | Additional Information |
|---|---|---|---|---|---|---|---|---|
| Osteoporosis | M80; M800; M8000; M8001; M8002; M8003; M8004; M8005; M8006; M8007; M8008; M8009; M805; M8050; M8051; M8052; M8053; M8054; M8055; M8056; M8057; M8058; M8059; M808; M8080; M8081; M8082; M8083; M8084; M8085; M8086; M8087; M8088; M8089; M809; M8090; M8091; M8092; M8093; M8094; M8095; M8096; M8097; M8098; M8099; M81; M810; M8100; M8101; M8102; M8103; M8104; M8105; M8106; M8107; M8108; M8109; M815; M8150; M8151; M8152; M8153; M8154; M8155; M8156; M8157; M8158; M8159; M816; M8160; M8161; M8162; M8163; M8164; M8165; M8166; M8167; M8168; M8169; M818; M8180; M8181; M8182;M8183; M8184; M8185; M8186; M8187; M8188; M8189; M819; M8190; M8191; M8192; M8193; M8194; M8195; M8196; M8197; M8198; M8199 | 7330; 73300; 73301; 73302; 73303; 73304; 73305; 73306; 73307; 73308; 73309 | | 1309 | | | Codes for osteoporosis | Control group excludes all individuals with osteopenia or with secondary forms of osteoporosis (including post-oophorectomy, disuse osteoporosis, drug-induced osteoporosis, or secondary to other diseases, e.g. multiple myelomatosis) |
| Myocardial Infarction and/or Coronary Revascularization | I21; I210; I211; I212; I213; I214; I219; I21X; I22; I220; I221; I228; I229; I23; I230; I231; I232; I233; I234; I235; I236; I238; I24; I240; I241; I248; I249; I252 | 410; 4109; 411; 4119; 412; 4129 | K40; K401; K402; K403; K404; K408; K409; K41; K411; K412; K413; K414; K418; K419; K42; K421; K422; K423; K424; K428; K429; K43; K431; K432; K433; K434; K438; K439; K44; K441; K442; K448; K449; K45; K451; K452; K453; K454; K455; K456; K458; K459; K46; K461; K462; K463; K464; K465; K468; K469; K49; K491; K492; K493; K494; K498; K499; K501; K75; K751; K752; K753; K754; K758; K759 | 1075 | 1070; 1095; 1523 | 1 | Codes for myocardial infarction, percutaneous transluminal coronary angioplasty and coronary artery bypass grafting. | Outcome definition as per Nelson et al (*HARD CAD*); Control group excludes all individuals with CHD and ICD-10 (I250; I253; I254) and ICD-9 codes (4141) pertaining to 'heart aneurysm' or 'atherosclerotic cardiovascular disease'. |
| Coronary Heart Disease | I20; I200; I201; I208; I209; I21; I210; I211; I212; I213; I214; I219; I21X; I22; I220; I221; I228; I229; I23; I230; I231; I232; I233; I234; I235; I236; I238; I24; I240; I241; I248; I249; I251; I252; I255; I256; I258; I259 | 410; 4109; 411; 4119; 412; 4129; 413; 4139; 4140; 4148; 4149 | K40; K401; K402; K403; K404; K408; K409; K41; K411; K412; K413; K414; K418; K419; K42; K421; K422; K423; K424; K428; K429; K43; K431; K432; K433; K434; K438; K439; K44; K441; K442; K448; K449; K45; K451; K452; K453; K454; K455; K456; K458; K459; K46; K461; K462; K463; K464; K465; K468; K469; K49; K491; K492; K493; K494; K498; K499; K501; K75; K751; K752; K753; K754; K758; K759 | 1074; 1075 | 1070; 1095; 1523 | 1; 2 | Codes for myocardial infarction, percutaneous transluminal coronary angioplasty, coronary artery bypass grafting, chronic ischemic heart disease and angina. | Outcome definition as per Nelson et al (*SOFT CAD*); Control group excludes all individuals with CHD and ICD-10 (I250; I253; I254) and ICD-9 codes (4141) pertaining to 'heart aneurysm' or 'atherosclerotic cardiovascular disease'. |
| Heart Failure | I50; I501; I509 | 428; 4280; 4281; 4289 | | 1076 | | | Codes for heart failure (including congestive heart failure and left heart failure). | |
| All Stroke | I60; I600; I601; I602; I603; I604; I605; I606; I607; I608; I609; I61; I610; I611; I612; I613; I614; I615; I616; I618; I619; I63; I630; I631; I632; I633; I634; I635; I636; I638; I639; I64 | 430; 4309; 431; 4319; 434; 4340; 4341; 4349; 436; 4369 | | 1081; 1086; 1491; 1583 | | 3 | Codes for subarachnoid and intracerebral haemorrhages and cerebral infarctions including cerebral thrombosis and embolism, and unspecified stroke. Does not included transient cerebral ischemia but includes acute but ill-defined cerebrovascular disease. | We used a common control group for 'All Stroke', 'Ischemic Stroke' and 'Hemorrhagic Stroke'. Participants were excluded from the stroke control group if they had ICD-10 (G45; G450; G451; G452; G453; G454; G458; G459) or ICD-9 (1082) codes pertaining to transient ischemic attacks. |
| Ischemic Stroke | I63; I630; I631; I632; I633; I634; I635; I636; I638; I639 | 434; 4340; 4341; 4349 | | 1583 | | | Codes for ischemic stroke, cerebral infarction, thrombosis or embolism. | We used a common control group for 'All Stroke', 'Ischemic Stroke' and 'Hemorrhagic Stroke'. Participants were excluded from the stroke control group if they had ICD-10 (G45; G450; G451; G452; G453; G454; G458; G459) or ICD-9 (1082) codes pertaining to transient ischemic attacks. |
| Hemorrhagic Stroke | I60; I600; I601; I602; I603; I604; I605; I606; I607; I608; I609; I61; I610; I611; I612; I613; I614; I615; I616; I618; I619 | 430; 4309; 431; 4319 | | 1086; 1491 | | | Codes for subarachnoid and intracerebral haemorrhages. | We used a common control group for 'All Stroke', 'Ischemic Stroke' and 'Hemorrhagic Stroke'. Participants were excluded from the stroke control group if they had ICD-10 (G45; G450; G451; G452; G453; G454; G458; G459) or ICD-9 (1082) codes pertaining to transient ischemic attacks. |
| Peripheral Vascular Disease | I702; I7020; I7021; I739; I742; I743; I744; I745 | 4439; 4442; 4402 | | 1067; 1087 | | | Codes for peripheral vascular disease, intermittent leg claudication, atherosclerosis and/or embolism and thrombosis of the extremities. | |
| Aortic Aneurysm | I711; I712; I713; I714; I715; I716; I718; I719; I790 | 441; 4410; 4411; 4412; 4413; 4414; 4415; 4416 | L18; L181; L182; L183; L184; L185; L186; L188; L189; L19; L191; L192; L193; L194; L195; L196; L198; L199; L254; L27; L271; L272; L273; L274; L275; L276; L278; L279; L28; L281; L282; L283; L284; L285; L286; L288; L289 | 1492; 1591 | 1104 | | Codes for aortic aneurysm (thoracic, abdominal and non-specified; including rupture, repair of aneurysm or stent insertion) | |
| Aortic Stenosis | I350, I352 | | | 1409 | | | Codes for aortic valve stenosis | |
| Chronic Kidney Disease | N18; N180; N181; N182; N183; N184; N185; N188; N189 | 585; 5859 | | | | | Codes for chronic renal failure or chronic kidney disease specifically | We excluded any cases of renal insufficiency from the control group, including ICD-9 (584; 5845; 5846; 5847; 5848; 5849; 585; 5859; 586; 5869), ICD-10 (N17; N170; N171; N172; N178; N179; N18; N180; N181; N182; N183; N184; N185; N188; N189; N19; I120; I131; I132; Z992), OPCS4 (L746; X40; X401; X402; X403; X404; X405; X406; X407; X408; X409; X41; X411; X412; X418; X419; X42; X421; X428; X429), or self-reported codes (1192; 1193; 1194; 1476; 1580; 1581; 1582) pertaining to non-specific renal insufficiency, acute kidney injury and/or non-specific dialysis. |

The following data fields were used for each diagnosis code type; ICD-10 codes: 41202 (main diagnosis), 41204 (secondary diagnoses), 40001 (primary cause of death), 40002 (secondary causes of death); ICD-9: 41203 (main diagnosis), 41205 (secondary diagnoses), 40013 (cancer register); OPCS-4 (operative procedures): 41200 (main code), 41210 (secondary code); self-reported at nurse's interview, non-cancer codes: 20002; self-reported at nurses's interview, operation codes: 20004; self-reported diagnosis from doctor: 6150 (for stroke and CHD). ICD, International Classification of Diseases; CHD, coronary artery disease; NI, nurse interview.

**Table S4. Definitions of outcomes analysed in Partners HealthCare Biobank.**

| Trait | Controls | Cases | Total | Definition |
|---|---|---|---|---|
| Fracture | 14868 | 4264 | | Cases include patients have any of the following code: [ICD-9] 800-804 (fracture of skull), 805-809 (fracture of spine and trunk), 810-819 (fracture of upper limb), and 820-829 (fracture of lower limb) and [ICD-10] S02 (fracture of skull and facial bones), S12 (fracture of cervical vertebra and other parts of neck), S22 (fracture of rib(s), sternum and thoracic spine), S32 (fracture of lumbar spine and pelvis), S42 (fracture of shoulder and upper arm), S52 (fracture of forearm), S62 (fracture at wrist and hand level), S72 (fracture of femur, S82 (fracture of lower leg, including ankle), S92 (fracture of foot and toe, except ankle). Controls include patients without any code for fracture mentioned above. |
| Osteoporosis | 16124 | 3008 | | As per eTable 3 (only ICD-9 and ICD-10 were used). |
| Coronary heart disease | 12439 | 6599 | | As per eTable 3 (only ICD-9 and ICD-10 were used). |
| Myocardial infarction and/or coronary revascularization | 12439 | 3288 | | As per eTable 3 (only ICD-9 and ICD-10 were used). |
| Type 2 diabetes | 16837 | 1951 | | EHR-based phenotyping algorithm with positive predictive value [PPV] = 0.9 for cases and negative predictive value [NPV] = 0.99 for controls. |
| Body mass index | | | 16154 | Median value for each patient from all available records. 5 individuals with BMI > 100 or < 10 were excluded. BMI were first regressed on sex, age at assessment, (age at assessment) squared, genotype batches and 15 principal components. The resulting residuals were inverse rank normalized. |
| HDL-cholesterol | | | 13032 | Median value for each patient from all available records. HDL-cholesterol were first regressed on sex, age at assessment, (age at assessment) squared, genotype batches and 15 principal components. The resulting residuals were inverse rank normalized. |
| LDL-cholesterol | | | 13032 | Median value for each patient from all available records. LDL-cholesterol were first regressed on sex, age at assessment, (age at assessment) squared, genotype batches and 15 principal components. The resulting residuals were inverse rank normalized. |
| Triglycerides | | | 13032 | Median value for each patient from all available records. Triglycerides were first regressed on sex, age at assessment, (age at assessment) squared, genotype batches and 15 principal components. The resulting residuals were inverse rank normalized. |
| Hypertension | 8917 | 8637 | | EHR-based phenotyping algorithm with positive predictive value [PPV] = 0.9 for cases and negative predictive value [NPV] = 0.99 for controls. |

#### Table S5. Definitions of outcomes analysed in China Kadoorie Biobank.

| Outcome | Case definition | Control definition | Analysis |
|---|---|---|---|
| Fracture | Self-reported fracture where baseline age minus age at diagnosis is <=5 years | No fracture (at any age), individual in random population. | Logistic regression stratified by region, adjusted for age, sex and first four regional PCs. |
| Osteoporosis | Combined endpoint: Self-reported osteoporosis plus M80 + M81<br>Self-reported and incident case also analysed separately. | Common controls: No self-reported or incident osteoporosis, individual in random population | Logistic regression stratified by region, adjusted for age, sex and first four regional PCs. |
| Coronary heart disease | CHD (I20-I25) plus revascularisation | Individual in random population, no self-reported CHD, no self-reported stroke or tia, no incident stroke (I60-I61, I63-I64), no death underlying from vascular disease (I00-I99). | Logistic regression stratified by region, adjusted for age, sex and first four regional PCs. |
| Myocardial infarction and/or coronary revascularisation | I21-I23 plus revascularisation | Individual in random population, no self-reported CHD, no self-reported stroke or tia, no incident stroke (I60-I61, I63-I64), no death underlying from vascular disease (I00-I99). | Logistic regression stratified by region, adjusted for age, sex and first four regional PCs. |
| Type 2 diabetes mellitus | Incident diabetes of any kind (E10:E14) plus self-reported/screen-detected where age of diagnosis is more than 30. | Individual in random population and no incident diabetes or self-reported/screen-detected diabetes (at any age). | Logistic regression stratified by region, adjusted for age, sex and first four regional PCs. |
| Hypertension | Self-reported hypertension or incident hypertension (I10) | No hypertension, individuals in random population subset | Logistic regression stratified by region, adjusted for age, sex and first four regional PCs. |
| Estimated bone mineral density | Apply exclusions separately in left and right foot:<br>- SOS: <=1,450 or >=1,700 m/s for males, <=1,455 or >=1,700 m/s for females<br>- BUA: <=27 or >=138 dB/MHz for males, <=22 or >=138 dB/MHz for females.<br>Calculate eBMD in each foot:<br>- eBMD = 0.002592 × (BUA + SOS) – 3.687<br>BMD Exclusions:<br>- eBMD: <=0.18 or >=1.06 g/cm2 for males; <=0.12 or >=1.025 g/cm2 for females.<br>Take mean of valid eBMD values | | Linear regression stratified by region, adjusted for age, sex and first four regional PCs. |
| BMI | | | Adjustment (stratified by region) for age, age2, sex and first four regional PCs. RINT of residuals (stratified by region). Linear regression (stratified by region) |
| WHR adjusted for BMI | | | Adjustment (stratified by region) for bmi, age, age2, sex and first four regional PCs. RINT of residuals (stratified by region). Linear regression (stratified by region) |
| HDL cholesterol | Exclude values with invalid flag. | | Adjustment (stratified by region) for age, age2, sex, ascertainment (x3) and first four regional PCs. RINT of residuals (stratified by region). Linear regression (stratified by region) |
| LDL cholesterol | Exclude values with invalid flag. | | Adjustment (stratified by region) for age, age2, sex, ascertainment (x3) and first four regional PCs. RINT of residuals (stratified by region). Linear regression (stratified by region) |
| Triglycerides | Exclude values with invalid flag. | | Adjustment (stratified by region) for age, age2, sex, ascertainment (x3) and first four regional PCs. RINT of residuals (stratified by region). Linear regression (stratified by region) |
| Systolic blood pressure | Mean SBP. Added 15mmHg for anyone reporting blood pressure medication (including ace-inhibitor, beta-blocker, diuretics and/or calcium channel blockers). | | Adjustment for age, sex, region, mean temperature (minimum value 5C, set any lower value to 5) and first four regional PCs. Linear regression of residuals (stratified by region) |
| Diastolic blood pressure | Mean DBP. Added 10mmHg for anyone reporting blood pressure medication (including ace-inhibitor, beta-blocker, diuretics and/or calcium channel blockers). | | Adjustment for age, sex, region, mean temperature (minimum value 5C, set any lower value to 5) and first four regional PCs. Linear regression of residuals (stratified by region) |

#### Table S6. Published phase III randomized controlled trials of romosozumab.

| Trial | Design | Year of Publication | PMID | Population | Arms | Duration | Intervention | | Comparator | | Total Randomized (Intervention / Comparator) | Primary outcome(s) |
|---|---|---|---|---|---|---|---|---|---|---|---|---|
| | | | | | | | 0-12 months | 13-24 months | 0-12 months | 13-24 months | | |
| FRAME | Placebo-controlled, double-blind | 2016 | 28892457 | Post-menopausal women with osteoporosis | 2 | 24 months | Romosozumab, 210mg monthly | Denosumab, 60mg, 6-monthly | Placebo | Denosumab, 60mg, 6-monthly | 7180 (3589 / 3591) | Cumulative incidences of new vertebral fracture at 12 months and at 24 months |
| ARCH | Active-controlled, double-blind | 2017 | 29424257 | Post-menopausal women with osteoporosis and previous fragility fracture | 2 | 24 months | Romosozumab, 210mg monthly | Alendronate, 70mg, weekly | Alendronate | Alendronate, 70mg, weekly | 4093 (2046 / 2047) | Cumulative incidence of new vertebral fracture at 24 months; cumulative incidence of clinical fracture |
| STRUCTURE | Active-controlled, open-label | 2017 | 28755782 | Postmenopausal women with osteoporosis transitioning from oral bisphosphonate therapy | 2 | 12 months | Romosozumab, 210mg monthly | NA | Teriparatide | NA | 436 (218 / 218) | Percentage change from baseline in total hip BMD at month 12. |
| BRIDGE | Placebo-controlled, double-blind | 2018 | 29931216 | Men with osteoporosis +- previous fragility fracture | 2 | 12 months | Romosozumab, 210mg monthly | NA | Placebo | NA | 245 (163 / 82) | Percentage change from baseline in lumbar spine BMD at month 12. |

PMID refers to PubMed identification number of the principal publication in which the trial was first reported.

**Table S7. Reported cardiovascular outcomes at 12 months from published phase III randomized controlled trials of romosozumab.**

| | BRIDGE | | ARCH | | | | FRAME | | | |
|---|---|---|---|---|---|---|---|---|---|---|
| | 12 months | | 12 months | | 24 months | | 12 months | | 24 months | |
| **Event** | **Romosozumab*** | **Placebo** | **Romosozumab*** | **Alendronate** | **Romosozumab to Alendronate** | **Alendronate to Alendronate** | **Romosozumab*** | **Placebo** | **Romosozumab to Denosumab** | **Placebo to Denosumab** |
| | N=163 | N=81 | N=2040 | N=2014 | N=2040 | N=2014 | N=3581 | N=3576 | N=3581 | N=3576 |
| | *number of patients (%)* | | | | | | | | | |
| Cardiac ischemic event | 3 (1.8%) | 0 (0%) | 16 (0.8%) | 6 (0.3%) | 30 (1.5%) | 20 (1%) | NA | NA | NA | NA |
| Cerebrovascular event | 3 (1.8%) | 1 (1.2%) | 16 (0.8%) | 7 (0.3%) | 45 (2.2%) | 27 (1.3%) | NA | NA | NA | NA |
| Serious cardiovascular event | 8 (4.9%) | 2 (2.5%) | 50 (2.5%) | 38 (1.9%) | 133 (6.5%) | 122 (6.1%) | 44 (1.2%) | 41 (1.1%) | 82 (2.3%) | 79 (2.2%) |

*Romosozumab 210mg monthly for 12 months. NA means events for specific outcome were not reported in publication.

**Table S8. Summary statistics of meta-analyses of cardiovascular events in randomized controlled trials of romosozumab.**

| Event | Romosozumab-arm | | Comparator-arm | | MH-method | | | Peto-method | | |
|---|---|---|---|---|---|---|---|---|---|---|
| | Events | Total | Events | Total | OR (95% CI) | P value | P-het | OR (95% CI) | P value | P-het |
| Cardiac Ischemic Events | 19 | 2,203 | 6 | 2,095 | 2.98 (1.18–7.55) | 0.02 | 0.78 | 2.63 (1.19–5.80) | 0.02 | 0.64 |
| Cerebrovascular event | 19 | 2,203 | 8 | 2,095 | 2.15 (0.94–4.92) | 0.07 | 0.74 | 2.06 (0.96–4.41) | 0.06 | 0.73 |
| Serious cardiovascular event | 102 | 5,784 | 81 | 5,671 | 1.21 (0.90–1.63) | 0.20 | 0.65 | 1.21 (0.90–1.62) | 0.20 | 0.66 |

MH, Mantel-Haenzsel meta-analysis; OR, odds ratio; CI, confidence interval; P-het refers to p value from Cochran’s Q test.

**Table S9. Scaled estimates for glycaemic traits from MAGIC consortium.**

| Trait | PMID | N | SNP | EA | OA | Units | Effect size | SE | P value |
|---|---|---|---|---|---|---|---|---|---|
| HbA1C | 28898252 | 123,665 | rs7220711 | G | A | % units | -0.009 | 0.02 | 0.69 |
| Fasting Glucose | NA | 68,784 | rs7220711 | G | A | mmol/l | 0.06 | 0.03 | 0.05 |
| Fasting Insulin | NA | 54,213 | rs7220711 | G | A | log(mmol/l) | -0.01 | 0.03 | 0.74 |

Estimates are scaled to match the effect of 210mg romosozumab monthly for 12 months on lumbar spine bone mineral density (0.09 g/cm2; see Methods) and aligned to the BMD-increasing alleles. PMID refers to PubMed identification number of the principal publication within which the data-source was first reported. HbA1c, glycated haemoglobin; SNP, single nucleotide polymorphism; EA, effect allele; OA, other allele

#### Table S10. Summary statistics of meta-analyses of scaled allelic estimates.

| Outcome/Trait | Figure with per-SNP estimates* | N estimates | Cases | Controls | Total | Scaled effect (95% CI) | Scaled OR (95% CI) | P value | P-heterogeneity |
|---|---|---|---|---|---|---|---|---|---|
| Osteoporosis | eFigure 8 | 6 | 15239 | 462099 | NA | NA | 0.43 (0.36 – 0.52) | 2.4E-18 | 0.003 |
| Any fracture | eFigure 8 | 6 | 53074 | 423955 | NA | NA | 0.59 (0.54 – 0.66) | 1.4E-24 | 0.85 |
| Myocardial Infarction and/or coronary revascularization | eFigure 10 | 8 | 69649 | 556342 | NA | NA | 1.18 (1.06 – 1.32) | 2.8E-03 | 0.12 |
| Coronary heart disease | eFigure 10 | 8 | 106329 | 551647 | NA | NA | 1.10 (1.00 – 1.20) | 0.04 | 0.05 |
| Upper limb fracture | eFigure 9 | 2 | 12823 | 379067 | NA | NA | 0.46 (0.38 – 0.56) | 6.1E-15 | 0.15 |
| Lower limb fracture | eFigure 9 | 2 | 9320 | 379067 | NA | NA | 0.62 (0.50 – 0.78) | 4.0E-05 | 0.62 |
| Vertebral fracture | eFigure 9 | 2 | 1016 | 379067 | NA | NA | 0.63 (0.32 – 1.26) | 0.19 | 0.42 |
| Other fracture | eFigure 9 | 2 | 23171 | 379067 | NA | NA | 0.64 (0.56 – 0.74) | 3.1E-09 | 0.58 |
| Type 2 diabetes mellitus | eFigure 11 | 4 | 76075 | 840843 | NA | NA | 1.15 (1.05 – 1.27) | 2.8E-03 | 0.49 |
| Hypertension | eFigure 11 | 4 | 235830 | 205207 | NA | NA | 1.12 (1.05 – 1.20) | 8.9E-04 | 0.2 |
| Atrial fibrillation | eFigure 11 | 2 | 60620 | 970216 | NA | NA | 1.05 (0.95 – 1.17) | 0.31 | 0.83 |
| Heart failure | eFigure 11 | 2 | 5840 | 417924 | NA | NA | 0.96 (0.72 – 1.27) | 0.77 | 0.98 |
| All stroke | eFigure 11 | 4 | 51398 | 818650 | NA | NA | 1.06 (0.94 – 1.19) | 0.37 | 0.7 |
| Ischemic stroke | eFigure 11 | 4 | 37581 | 816959 | NA | NA | 1.09 (0.94 – 1.25) | 0.25 | 0.32 |
| Hemorrhagic stroke | eFigure 11 | 2 | 1929 | 410848 | NA | NA | 1.18 (0.72 – 1.91) | 0.51 | 0.3 |
| Peripheral vascular disease | eFigure 11 | 2 | 4209 | 419555 | NA | NA | 1.04 (0.75 – 1.44) | 0.82 | 0.98 |
| Aortic aneurysm | eFigure 11 | 2 | 1527 | 422237 | NA | NA | 1.15 (0.66 – 1.99) | 0.62 | 0.23 |
| Aortic stenosis | eFigure 11 | 2 | 1954 | 421810 | NA | NA | 0.74 (0.45 – 1.20) | 0.22 | 0.9 |
| Chronic kidney disease | eFigure 11 | 3 | 16988 | 518923 | NA | NA | 0.88 (0.70 – 1.10) | 0.26 | 0.46 |
| Systolic blood pressure | eFigure 12 | 4 | NA | NA | 459357 | 1.33 (0.76 – 1.91) mmHg | NA | 5.9E-06 | 0.5 |
| Diastolic blood pressure | eFigure 12 | 2 | NA | NA | 423353 | -0.01 (-0.37 – 0.35) mmHg | NA | 0.95 | 0.12 |
| WHR adj. for BMI | eFigure 13 | 3 | NA | NA | 552964 | 0.05 (0.02 – 0.08) standard deviations | NA | 8.5E-04 | 0.54 |
| BMI | eFigure 13 | 5 | NA | NA | 660747 | 0.02 (-0.01 – 0.05) standard deviations | NA | 0.14 | 0.56 |
| Triglycerides | eFigure 13 | 3 | NA | NA | 102501 | 0.11 (0.01 – 0.20) standard deviations | NA | 0.02 | 0.92 |
| LDL cholesterol | eFigure 13 | 3 | NA | NA | 101450 | -0.03 (-0.13 – 0.07) standard deviations | NA | 0.57 | 0.92 |
| HDL cholesterol | eFigure 13 | 3 | NA | NA | 105836 | -0.09 (-0.18 – 0.01) standard deviations | NA | 0.07 | 0.91 |
| eGFR | eFigure 14 | 2 | NA | NA | 218678 | 0.01 (-0.01 – 0.02) log(ml/min/1.73 m2) | NA | 0.40 | 0.97 |

*Refers to figure where individual estimates are presented. N estimates refers to number of individual estimates meta-analysed per outcome/trait. P-heterogeneity refers to p value from Cochran's Q test, performed for each meta-analysis. CI, confidence interval; GFR, glomerular filtration rate; mmHg, millimeters of mercury; OR, odds ratio.

**Table S11. Per-allele estimates for rs7209826 in China Kadoorie Biobank.**

| Trait | SNP | Beta/log(OR) | SE of beta/log(OR) | p-value | Cases | Controls | Total |
|---|---|---|---|---|---|---|---|
| eBMD (left foot) | rs7209826 | 0.0021 | 0.00108 | 0.0523 | NA | NA | 21130 |
| eBMD (mean) | rs7209826 | 0.00157 | 0.00106 | 0.1373 | NA | NA | 21547 |
| eBMD (right foot) | rs7209826 | 0.00119 | 0.00108 | 0.2722 | NA | NA | 21079 |
| Osteoporosis (combined) | rs7209826 | 0.0176 | 0.0696 | 0.8 | 537 | 19221 | 19758 |
| Osteoporosis (incident) | rs7209826 | 0.2415 | 0.1669 | 0.1479 | 86 | 8031 | 8117 |
| Osteoporosis (self-reported) | rs7209826 | -0.00814 | 0.0789 | 0.9178 | 420 | 17354 | 17774 |
| Fracture | rs7209826 | -0.0298 | 0.0374 | 0.4246 | 2000 | 17899 | 19899 |
| Myocardial Infarction | rs7209826 | -0.0125 | 0.0246 | 0.6117 | 4901 | 60921 | 65822 |
| All CHD | rs7209826 | 0.00803 | 0.0162 | 0.6193 | 14948 | 57638 | 72586 |
| Type 2 Diabetes | rs7209826 | 0.00417 | 0.0179 | 0.8156 | 9158 | 66835 | 75993 |
| Hypertension | rs7209826 | -0.0201 | 0.0134 | 0.1327 | 22185 | 59361 | 81546 |
| HDL cholesterol | rs7209826 | 0.0147 | 0.0118 | 0.2154 | NA | NA | 17285 |
| LDL cholesterol | rs7209826 | 0.00616 | 0.0118 | 0.6027 | NA | NA | 17285 |
| Triglycerides | rs7209826 | -0.0101 | 0.0118 | 0.3955 | NA | NA | 17285 |
| BMI | rs7209826 | 0.000287 | 0.00576 | 0.9603 | NA | NA | 72795 |
| WHR adjusted for BMI | rs7209826 | -0.00184 | 0.00576 | 0.7494 | NA | NA | 72795 |
| Diastolic blood pressure | rs7209826 | -0.027 | 0.0649 | 0.6772 | NA | NA | 72796 |
| Systolic blood pressure | rs7209826 | -0.0179 | 0.1151 | 0.8762 | NA | NA | 72796 |

**Table S12. Reported fracture outcomes at 12 and 24 months from published phase III randomized controlled trials of romosozumab.**

| Outcome & Effect Estimate | ARCH | | FRAME | |
|---|---|---|---|---|
| | 12 months | | 12 months | |
| | Romosozumab | Alendronate | Romosozumab | Placebo |
| | N=2046 | N=2047 | N=3589 | N=3591 |
| | *number of patients (%)* | | | |
| Clinical fracture | 79 (3.9%) | 110 (5.4%) | 58 (1.6%) | 90 (2.5%) |
| Hazard Ratio (95% CI) | 0.72 (0.54, 0.96) | | 0.64 (0.46–0.89) | |

Clinical fracture is a composite of nonvertebral fracture and symptomatic vertebral fracture. CI, confidence interval.
